# Host specific sensing of coronaviruses and picornaviruses by the CARD8 inflammasome

**DOI:** 10.1101/2022.09.21.508960

**Authors:** Brian V. Tsu, Rimjhim Agarwal, Nandan S. Gokhale, Jessie Kulsuptrakul, Andrew P. Ryan, Lennice K. Castro, Christopher M. Beierschmitt, Elizabeth A. Turcotte, Elizabeth J. Fay, Russell E. Vance, Jennifer L. Hyde, Ram Savan, Patrick S. Mitchell, Matthew D. Daugherty

## Abstract

Hosts have evolved diverse strategies to respond to microbial infections, including the detection of pathogen-encoded proteases by inflammasome-forming sensors such as NLRP1 and CARD8. Here, we find that the 3CL protease (3CL^pro^) encoded by diverse coronaviruses, including SARS-CoV-2, cleaves a rapidly evolving region of human CARD8 and activates a robust inflammasome response. CARD8 is required for cell death and the release of pro-inflammatory cytokines during SARS-CoV-2 infection. We further find that natural variation alters CARD8 sensing of 3CL^pro^, including 3CL^pro^-mediated antagonism rather than activation of megabat CARD8. Likewise, we find that a single nucleotide polymorphism (SNP) in humans reduces CARD8’s ability to sense coronavirus 3CL^pros^, and instead enables sensing of 3C proteases (3C^pro^) from select picornaviruses. Our findings demonstrate that CARD8 is a broad sensor of viral protease activities and suggests that CARD8 diversity contributes to inter- and intra-species variation in inflammasome-mediated viral sensing and immunopathology.

## Introduction

Effector-triggered immunity (ETI) is a host defense strategy by which innate immune sensors recognize pathogens via the detection of pathogen-specific activities (*1–5*). A subset of eukaryotic ETI sensors form inflammasomes—large, intracellular immune complexes that activates a pro-inflammatory caspase, predominantly caspase-1 (CASP1), to initiate inflammatory signaling via interleukin (IL)-1β and IL-18 and pyroptotic cell death through cleavage of the pore-forming protein Gasdermin D (GSDMD) (*6–8*). In humans, the activity of viral proteases can be sensed by the inflammasome-forming sensors CARD8 and NLRP1 (*9–12*). CARD8 is comprised of a disordered N-terminal region and a C-terminal function-to-fund domain (FIIND) and caspase activation and recruitment domain (CARD). The FIIND undergoes self-cleavage resulting in a bipartite sensor, with the disordered N-terminus acting as a ‘tripwire’ for viral proteases. For instance, proteolytic cleavage of CARD8 by the HIV-1 protease (HIV-1^pro^) leads to proteasome-dependent ‘functional degradation’ (*13, 14*) of the cleaved N-terminus and release of the bioactive CARD-containing C-terminus, which is sufficient for inflammasome assembly and activation (*12*). However, the extent to which CARD8 has evolved to sense other viral proteases and functions as an innate immune sensor of viral infection has been unclear.

### The SARS-CoV-2 3CL^pro^ activates the human CARD8 inflammasome via proteolysis within the disordered N-terminus

To determine if the CARD8 inflammasome can sense coronavirus infection by mimicking sites of viral polyprotein cleavage, we expanded our previous bioinformatic approach (*11*) to generate a predictive model for the *Coronaviridae* main protease (3CL^pro^, also known as nsp5 or M protease), which has been shown to cleave host proteins in addition to the coronavirus polyprotein (*15–19*). The resulting 3CL^pro^ cleavage motif, XXΦQ[G/A/S]XXX (where Φ denotes a hydrophobic residue and X denotes any amino acid) (**Fig 1A, S1 Fig, S1 and S2 Tables**) is broadly consistent with previous studies (*20–23*), and allowed us to predict two putative 3CL^pro^ cleavage sites within the N-terminus of human CARD8 (**Fig 1A**).

**Fig 1.**
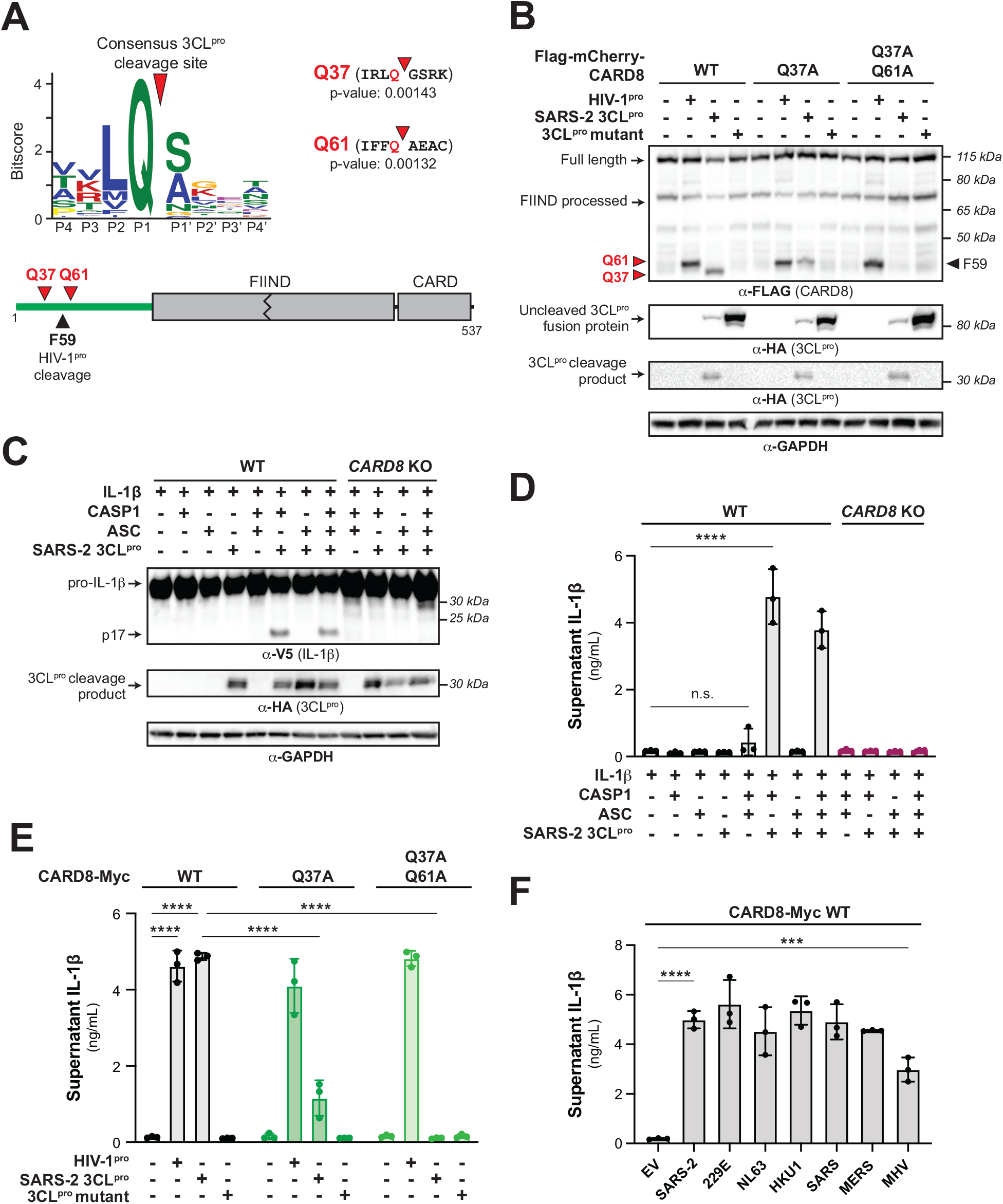
3CL^pro^ from SARS-CoV-2 and other coronaviruses cleaves and activates the human CARD8 inflammasome. (A) A consensus betacoronavirus 3CL^pro^ cleavage motif (*upper* panel, Fig S1 and Methods) was used to predict two 3CL^pro^ cleavage sites (red triangles) within the disordered ‘tripwire’ N-terminus of human CARD8 (*lower* panel, green) near the described site of HIV-1^pro^ cleavage (black triangle). Flanking residues and p-values of prediction for each site (Q37 and Q61) are shown. (B) HEK293T cells were transfected with the indicated CARD8 construct in the presence (‘+’) or absence (‘-’) of indicated proteases. Active (SARS-2 3CL^pro^) or catalytically inactive (3CL^pro^ mutant) protease from SARS-CoV-2 was expressed as an HA-tagged fusion construct (Fig S2). HIV-1^pro^ was expressed from an untagged gag-pol construct. Triangles are as described in (A). (C-D) WT or *CARD8* KO HEK293T cells were transfected with (‘+’) or without (‘-’) indicated constructs. Inflammasome activation was monitored by immunoblotting for mature IL-1β (p17) (C) or measuring culture supernatant levels of bioactive IL-1β using IL1R-expressing reporter cells (D). (E-F) *CARD8* KO HEK293T cells were co-transfected with the indicated CARD8 and protease constructs and supernatant levels of bioactive IL-1β were measured by IL1R reporter assay. 3CL^pros^ from the following viruses were used: HCoV-229E (229E), HcoV-NL63 (NL63), HcoV-HKU1 (HKU1), SARS-CoV (SARS), MERS-CoV (MERS), mouse hepatitis virus (MHV). (D-F) Individual values (n=3), averages, and standard deviations shown are representative of experiments performed in triplicate. Data were analyzed using two-way ANOVA with Šidák’s post-test (D-E) or one-way ANOVA with Tukey’s post-test (F). *** = p<0.001, **** = p<0.0001, n.s. = not significant.

To determine if human CARD8 is cleaved by coronavirus 3CL^pro^, we co-expressed an N-terminal 3xFlag/mCherry-tagged isoform of human CARD8 (wild-type (WT)) with 3CL^pro^ from SARS-CoV-2 in HEK293T cells (**Fig 1B** **and S2 Fig**). We used HIV-1^pro^ as a positive control since it had previously been shown to cleave CARD8 between residues F59-F60 (site F59 in **Fig 1, A and B**). We observed a ∼34kDa CARD8 product in the presence of SARS-CoV-2 3CL^pro^ but not the catalytically inactive C145A 3CL^pro^ mutant. The ∼34kDa product is predicted to result from cleavage at site Q37 (**Fig 1A**), which migrated slightly below the cleavage product of HIV-1^pro^. Mutating the putative P1 residue in the Q37 site (CARD8 Q37A) eliminated the 34kDa product, confirming SARS-CoV-2 3CL^pro^ cleavage at this site. The CARD8 Q37A mutant also revealed a cryptic ∼37kDa 3CL^pro^-dependent product, which matches cleavage at the predicted site Q61 (**Fig 1A** **and S2 Fig**). The CARD8 Q37A Q61A mutant was completely insensitive to cleavage by SARS-CoV-2 3CL^pro^ (**Fig 1B**), whereas cleavage by the HIV-1^pro^ was unperturbed by either mutant. Thus, CARD8 can be cleaved by SARS-CoV-2 3CL^pro^ at amino acid sequences that mimic the coronavirus polyprotein cleavage site.

For both NLRP1 and CARD8, N-terminal proteolytic cleavage can activate CASP1 in a reconstituted inflammasome assay (*10-12, 14, 24*). Consistent with the prior observation that CARD8 is endogenously expressed in some HEK293T cell lines (*12*), transfection of our HEK293T cells with only CASP1, pro-IL-1β, and SARS-CoV-2 3CL^pro^ resulted in robust CASP1-dependent processing of pro-IL-1β to mature bioactive IL-1β (p17) as measured by immunoblot or IL-1β reporter assay (see **Methods**) (**Fig 1, C and D**). Inflammasome activation was not observed in cells in *CARD8* knock out (KO) cells.

For both NLRP1 and CARD8, N-terminal proteolytic cleavage can activate CASP1 in a reconstituted inflammasome assay (*10-12, 14, 24*). Validating a prior observation that CARD8 is endogenously expressed in some HEK293T cell lines (*12*), transfection of our HEK293T cells with CASP1, pro-IL-1β, and SARS-CoV-2 3CL^pro^ resulted in robust CASP1-dependent processing of pro-IL-1β to mature bioactive IL-1β (p17) as measured by immunoblot or IL-1β reporter assay (**Fig 1, C and D**). We found that inflammasome activation was CARD8-dependent since we did not observe CASP1-processed IL-1β in HEK293T *CARD8* knock out (KO) cells. To confirm that SARS-CoV-2 3CL^pro^ cleavage of CARD8 is responsible for inflammasome activation, we complemented *CARD8* KO cells with WT CARD8 or cleavage site mutants. Complementation with WT CARD8 rescued both HIV-1^pro^ and 3CL^pro^-induced inflammasome activation (**Fig 1E** **and S3 Fig**). In contrast, whereas CARD8 cleavage site mutants had no effect on HIV-1^pro^-induced inflammasome activation, 3CL^pro^-induced inflammasome activation was reduced or abolished in *CARD8* KO cells complemented with CARD8 Q37A or CARD8 Q37A Q61A, respectively (**Fig 1E** **and S3 Fig)**. These results validate that 3CL^pro^ site-specific cleavage is required for CARD8 inflammasome activation. As expected, 3CL^pro^ inflammasome activation did not occur in *CARD8* KO cells complemented with the CARD8 FIIND auto-processing mutant S297A mutant (**S4 Fig**) (*12, 25, 26*). Taken together, our results indicate that human CARD8 senses the proteolytic activity of the SARS-CoV-2 3CL^pro^, which drives inflammasome activation via functional degradation.

To test if proteases from other coronaviruses also cleave CARD8, we cloned 3CL^pros^ from other human-relevant beta-coronaviruses (SARS-CoV (SARS), MERS-CoV (MERS), HCoV-HKU1 (HKU1), two human alpha-coronaviruses (HCoV-229E (229E) and HCoV-NL63 (NL63), and the mouse beta-coronavirus murine hepatitis virus (MHV) (**S5 Fig**). Consistent with their structural and cleavage motif similarity (*27*), we found that every tested 3CL^pro^ was able to cleave and activate CARD8 in a site-specific manner (**Fig 1F** **and S6 Fig**). Thus, effector-triggered immunity by human CARD8 is a conserved pathway for sensing of both endemic and pandemic human coronaviruses.

### Coronavirus infection activates the CARD8 inflammasome

We next evaluated the consequences of CARD8 inflammasome activation by infecting the human monocyte-like cell line THP-1 with HCoV-229E. We found that HCoV-229E infected WT but not two independently-derived *CARD8* KO THP-1 cell lines underwent significant cell death (**Fig 2A**) and release of IL-1β (**Fig 2B**), a result similar to treatment with Val-boroPro (VbP), which specifically activates the CARD8 inflammasome in myeloid and lymphoid lineages (*26, 28*) (**S7 Fig**). SARS-CoV-2 infection (*29, 30*) or uptake via FcyR-mediated antibody-dependent enhancement (*31*) by monocytes or macrophages induces inflammasome activation. To determine if CARD8 senses and responds to SARS-CoV-2 infection in THP-1 cells, we engineered WT or *CARD8* KO THP-1 cells to express ACE2 and TMPRSS2 (THP-1^A+T^). As with HCoV-229E, we found that SARS-CoV-2 infection of THP-1^A+T^ WT but not *CARD8* KO cells induced both cell death and IL-1β release (**Fig 2, C and D**). Our results demonstrate that CARD8 is a bona fide innate immune sensor of viral infection.

**Fig 2.**
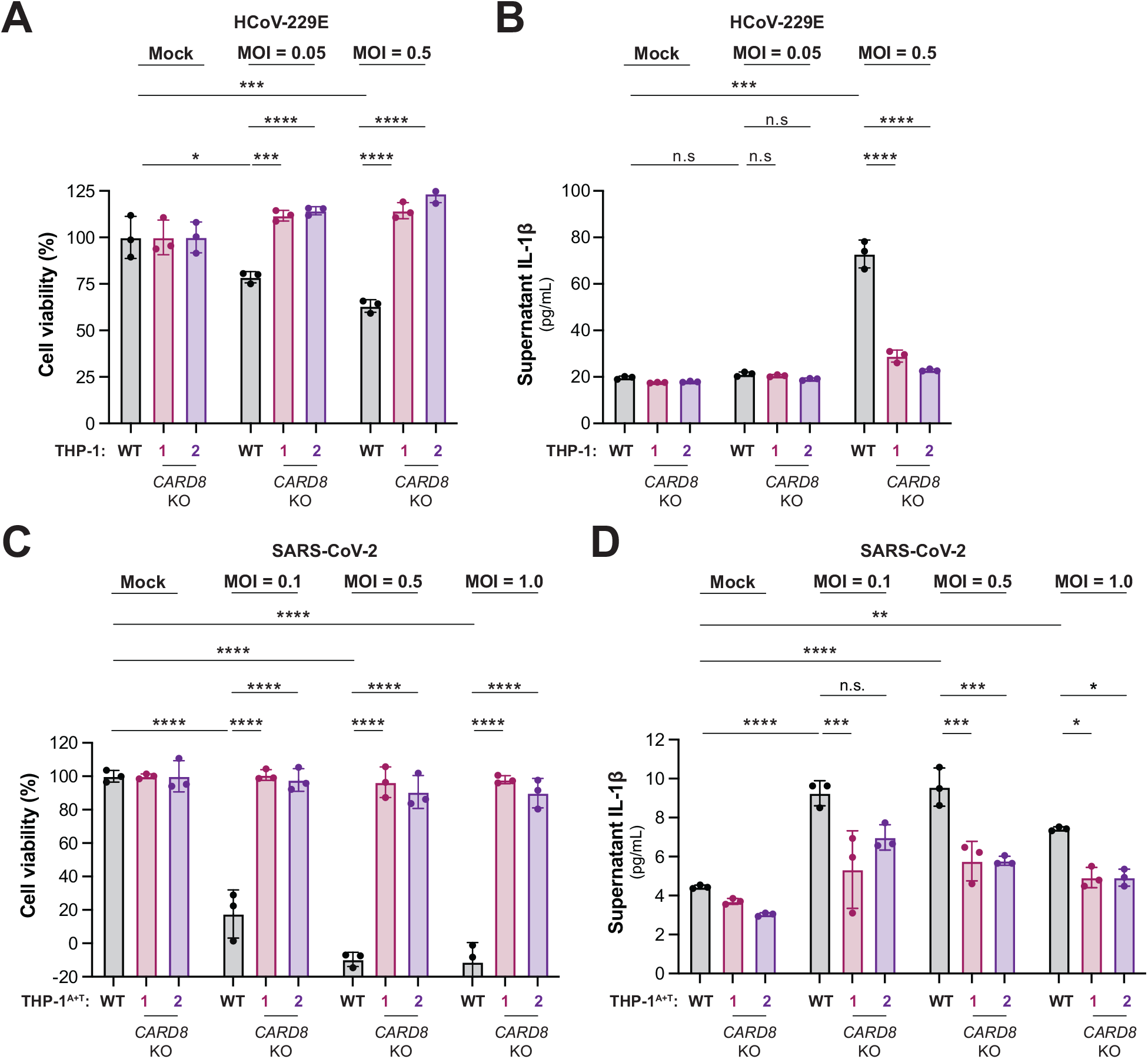
Coronavirus infection activates the CARD8 inflammasome in THP-1 cells. WT, *CARD8* KO1, and *CARD8* KO2 THP-1 cells (A-B) or THP-1 cells expressing ACE2 and TMPRSS2 (THP-1^T+A^) (C-D) were primed with 0.5 µg/mL Pam3CSK4 for 6h, followed by infection with the coronaviruses hCoV-229E (A-B) or SARS-CoV-2 (SARS-2) (C-D) at the indicated multiplicity of infection (MOI). 48h post-infection, cell viability (A, C) was measured using the Cell Titer Glo assay and IL-1β levels were measured using the IL1R reporter assay (B, D) as in Fig 1D. Data presented are representative of experiments performed at least twice. Data were analyzed using two-way ANOVA with Šidák’s post-test: * = p<0.05, ** = p<0.01, *** = p<0.001, **** = p<0.0001, n.s. = non-significant.

### Inter- and intra-host diversity in CARD8 impacts inflammatory responses to coronavirus proteases

Host-virus interactions, including those between viral proteases and host cleavage targets, are often engaged in evolutionary arms races that shape the specificity of host-virus interactions (*15, 32–34*). Indeed, we have previously shown that the CARD8 homolog and viral protease sensor, NLRP1, has been duplicated and recurrently lost across mammalian evolution. *NLRP1* also has strong signatures of positive selection in an N-terminal region of the protein that is cleaved by pathogen-encoded proteases, which we refer to as the ‘tripwire’ region due to its role in virus detection and subsequent inflammasome activation (*11, 24*). We thus predicted that *CARD8* may have a similarly dynamic evolutionary history, and that host inter- and intraspecies variation would underlie differences in CARD8 cleavage and inflammasome activation by coronavirus 3CL^pros^.

We first found that both *CARD8* and *NLRP1* are each present in only certain mammalian lineages, consistent with their dynamic roles in host defense as opposed to dedicated housekeeping functions (**Fig 3A**). For instance, we found that within the order *Chiroptera* (bats), microbats retain a *NLRP1* ortholog but have lost *CARD8*, whereas megabats have lost *NLRP1* and only encode *CARD8*. Because bats serve as main reservoir hosts of emerging coronaviruses (*35, 36*), but is missing from microbats, we tested if the CARD8 inflammasome could serve as a sensor for 3CL^pro^ in the megabat species *Rousettus aegyptiacus*. Unlike human CARD8, the *R. aegyptiacus* CARD8 (CARD8*_Ra_*) N-terminus lacks sites Q37 and Q61 and was not cleaved by SARS-CoV-2 3CL^pro^. We did, however, observe a cleavage product that matched a predicted 3CL^pro^ site at Q349 in CARD8*_Ra_* downstream of the FIIND auto-processing site in the inflammasome-forming C-terminus (**Fig 3, A and B**). Interestingly, megabats are the only mammals with a serine in the P1’ position of the cleavage site (**Fig 3A** **and S8 Fig)**, which is preferred for cleavage based on our computational model (**Fig 1A**). Indeed, a threonine in this position (S350T), which is found in human CARD8 as well as most other mammals, prevents cleavage of CARD8*_Ra_* by SARS-CoV-2 3CL^pro^ (**Fig 3C** **and S8 Fig**). We next wished to test the functional effect of 3CL^pro^ on the megabat CARD8 inflammasome. First, we determined if functional degradation could activate megabat CARD8 by inserting a TEV^pro^ cleavage site into the N-terminus of CARD8*_Ra_*, permitting TEV^pro^ cleavage of CARD8*_Ra_*-TEV but not WT CARD8*_Ra_*. When co-expressed with CASP1 and pro-IL-1β from *R. aegyptiacus*, TEV^pro^ cleavage of CARD8*_Ra_*-TEV resulted in inflammasome activation, indicating that we can reconstitute the *R. aegyptiacus* CARD8 inflammasome in human cells (**Fig 3D**). Using this reconstitution system, we found that SARS-CoV-2 3CL^pro^ does not activate, and in fact antagonizes TEV-mediated CARD8*_Ra_* inflammasome activation (**Fig 3D**), similar to our previous observations of viral proteases that antagonize the activation of the NLRP1 inflammasome (*11*). We further found that all 3CL^pros^ that we tested can instead prevent TEV^pro^-mediated activation of the CARD8*_Ra_* inflammasome (**Fig 3E**). Together with the loss of CARD8 from many bat species, these data provide a putative mechanism of disease tolerance that protects bats from immunopathogenic effects of inflammasome activation.

**Fig 3.**
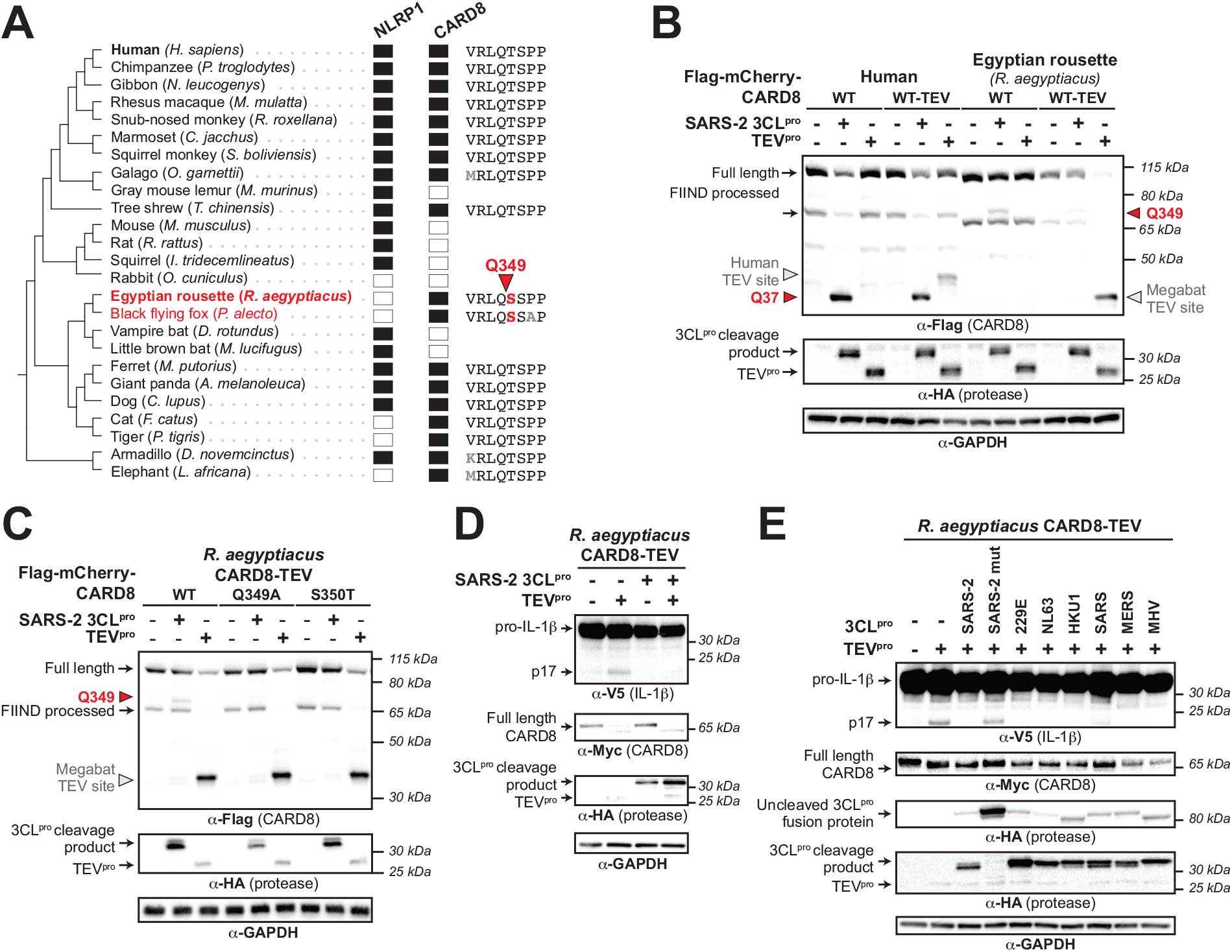
Megabat CARD8 is antagonized rather than activated by coronavirus 3CL^pro^. (A) Presence (filled rectangle) or absence (empty rectangle) of predicted *NLRP1* or *CARD8* orthologs in the indicated mammalian species. To the left is a species phylogeny. Megabat species are indicated in red. To the right is an alignment of a predicted 3CL^pro^ cleavage site in *Rousettus aegyptiacus* CARD8 (red triangle indicates site and number indicates residue position). (B) Human CARD8 or *R. aegyptiacus* CARD8 was co-transfected with either SARS-CoV-2 (SARS-2) 3CL^pro^ or protease from tobacco etch virus (TEV^pro^). For human or *R. aegyptiacus* CARD8 constructs labeled “WT-TEV”, a TEV^pro^ site was introduced into the N-terminus. The red triangles and amino acid number indicates the sites of 3CL^pro^ cleavage in human and *R. aegyptiacus* CARD8. The gray triangles indicate the sites of TEV protease cleavage within each CARD8 WT-TEV. (C) Mapping of the 3CL^pro^ site within *R. aegyptiacus* CARD8 was performed by transfecting the indicated point mutants with SARS-CoV-2 (SARS-2) 3CL^pro^ or TEV^pro^. 3CL^pro^ and TEV^pro^ sites are marked by triangles as in (B). (D) *CARD8* KO HEK293T cells were co-transfected with *R. aegyptiacus* IL-1β and CASP1, along with the indicated CARD8 and protease constructs. Presence of mature IL-1 β (p17) upon TEV^pro^ addition indicates successful reconstitution of the *R. aegyptiacus* CARD8 inflammasome, whereas absence of p17 upon SARS-2 3CL^pro^ indicates antagonism of the *R. aegyptiacus* CARD8 inflammasome. (E) *R. aegyptiacus* CARD8 inflammasome activation assays were performed as in (D) with the indicated 3CL^pro^ constructs.

We next focused on the evolution of human *CARD8*. Supporting a previous genome-wide study (*37*), we found evidence that *CARD8* has evolved under recurrent positive selection in hominoids and Old World monkeys, which we find is primarily driven by the N-terminal region of the protein (**Fig 4A**). Codon-based analyses also show that positively selected sites are predominantly found in the N-terminus, including a codon at position 60 that lies in the HIV-1^pro^ site and the secondary coronavirus 3CL^pro^ site (**Fig 4B, S9 Fig, and S5 Table)**. We thus infer that, like NLRP1, the CARD8 disordered N-terminus is a molecular ‘tripwire’ that is rapidly evolving to mimic viral polyprotein sites and sense diverse viral proteases. We further analyzed the human population for non-synonymous SNPs in *CARD8* (**Fig 4B**). Within the N-terminus, we found several high frequency human SNPs, including a S39P variant that resides at the P2’ position within the 3CL^pro^ cleavage site and is present in >20% of all sampled African and African American individuals (GnomAD v3.1.2 (*38*)) (**S9 Fig and S5 and S6 Tables**). Strikingly, while CARD8 S39 and P39 variants are similarly cleaved and activated by HIV-1^pro^, the CARD8 P39 variant exhibits reduced sensitivity to cleavage and activation by coronavirus 3CL^pros^ (**Fig 4, C and D**, and **S10 Fig**). This is reinforced by our observation that a proline in the P2’ position is never found in the >10,000 polyprotein cleavage sites we sampled from beta-CoVs (**S1 Table**). In contrast, another human SNP (R40W) found in approximately 1 in every 2000 alleles (**S5 Table**), does not detectably affect CARD8 cleavage in our assays (**Fig 4, C and D**). These data suggest that standing genetic variation in human CARD8 underlies differential sensing and inflammasome responses to coronavirus infection.

**Fig 4.**
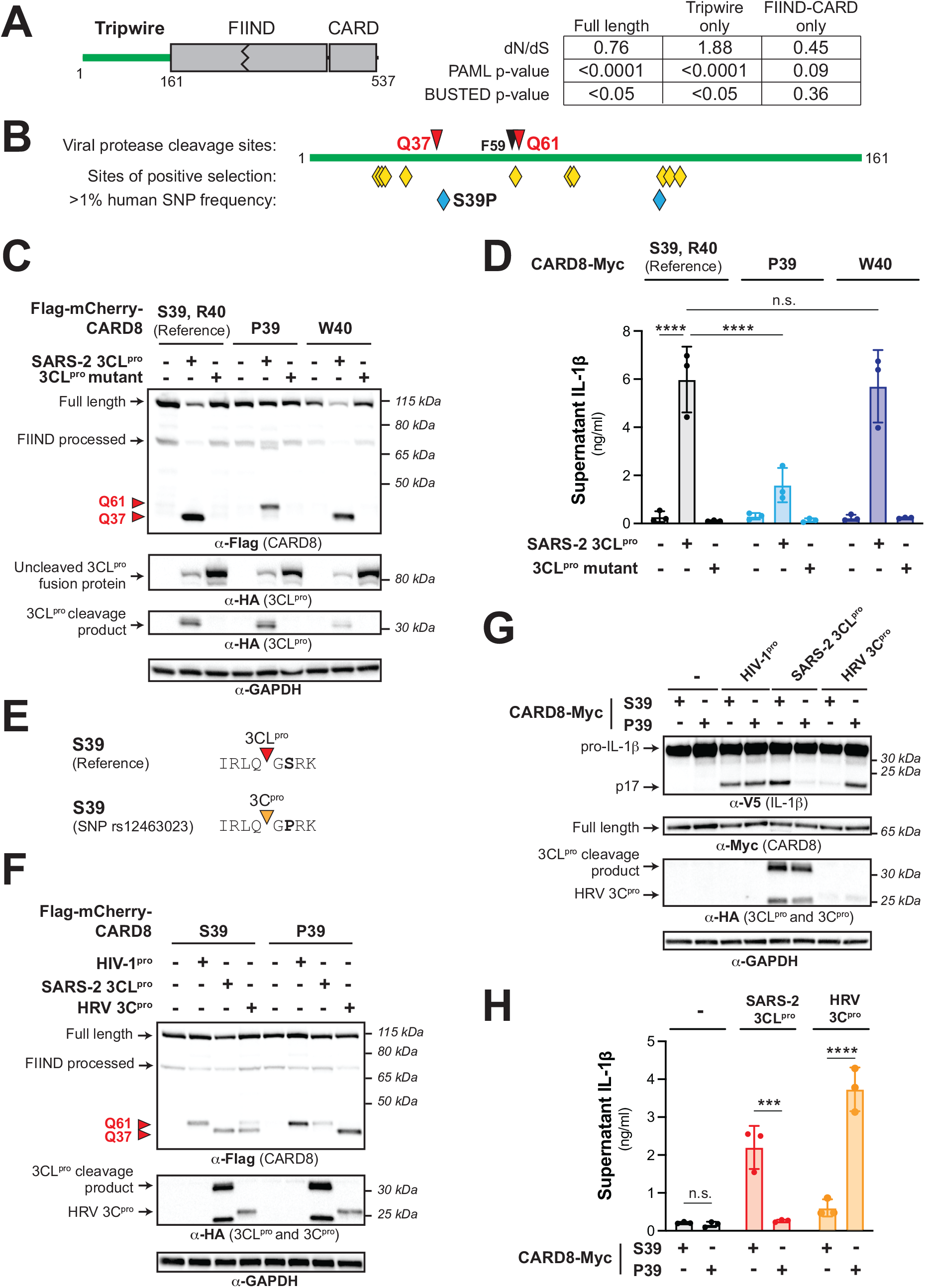
Human polymorphism in CARD8 reduces sensing of coronavirus 3CL^pro^ while increasing sensing of select picornavirus 3C^pros^. (A) Evolutionary analyses of positive selection were performed on full length *CARD8* (encoding residues 1-537), the disordered N-terminal ‘tripwire’ region (encoding residues 1-161), and the FIIND-CARD region (encoding residues 162-537). P-values from PAML and BUSTED analyses are shown, along with the dN/dS value obtained from PAML. (B) Schematic of the CARD8 ‘tripwire’ region. Red and black triangles and amino acid numbers indicate sites of 3CL^pro^ and HIV-1^pro^ cleavage respectively. Yellow diamonds indicate codons predicted to be evolving under positive selection by at least one evolutionary analysis (table S4). Blue diamonds indicate high frequency (>1% allele frequency) non-synonymous single nucleotide polymorphisms (SNPs) in humans (table S5 and S6). The position of the S39P substitution that results from SNP rs12463023 is shown. (C-D) Reference human CARD8 (S39, R40) or human CARD8 variants (P39 or W40) were co-expressed with the indicated protease construct and assayed for 3CL^pro^-mediated cleavage (C) or CARD8 inflammasome activation (D). (E) Amino acid sequence surrounding the Q37 cleavage site for S39 (reference) and P39 variants of human CARD8. Triangles mark sites of cleavage by the indicated viral protease (F-H). Human CARD8 S39 or CARD8 P39 were transfected with the indicated proteases and assayed for 3CL^pro^-mediated cleavage (F) or CARD8 inflammasome-mediated maturation of IL-1β (G) or the release of bioactive IL-1β (H). HRV 3C^pro^ = human rhinovirus 3C^pro^. Individual values (n=3), averages, and standard deviations shown are representative of experiments performed in triplicate. Data were analyzed using two-way ANOVA with Šidák’s post-test. *** = p<0.001, **** = p<0.0001, n.s. = not significant.

### A human SNP confers a specificity switch for CARD8 sensing of coronavirus 3CL^pro^ and human rhinovirus 3C^pro^

Finally, we considered if the CARD8 P39 variant, in addition to affecting sensing of coronavirus 3CL^pros^, also alters recognition of other pathogens. We noticed that although a P2’ proline is disfavored in our model for 3CL^pro^ cleavage, a P2’ proline is strongly preferred in our model for enterovirus 3C^pro^ cleavage (*11*). Indeed, our 3C^pro^ motif search (*11*) identified a cleavage site in CARD8 P39 but not CARD8 S39 (**Fig 4E** **and S11 Fig**). Validating these bioinformatic predictions, cleavage assays with the 3C^pro^ from the respiratory picornavirus, human rhinovirus (HRV), revealed that HRV 3C^pro^ cleavage of human CARD8 at site Q37 is considerably more pronounced for the P39 variant than the S39 variant (**Fig 4F**). Likewise, we found that inflammasome activation by HRV 3C^pro^ was nearly absent in HEK293T *CARD8* KO cells complemented with CARD8 S39, whereas we observed robust inflammasome activation in cells complemented with CARD8 P39 – the opposite sensitivity observed for SARS-CoV-2 3CL^pro^ (**Fig 4, G and H**). Finally, we found that CARD8 variation also impacted cleavage by other picornavirus 3C^pros^. For example, 3C^pros^ from enterovirus D68 (EV68) and poliovirus (PV1) were better sensed by CARD8 P39, whereas the Aichi virus 3C^pro^ was better sensed by the S39 CARD8 variant (**S12 Fig**). Thus, a single amino acid change in CARD8 functions as a viral-specificity switch, underscoring the importance of pathogen-driven evolution in shaping inflammasome responses.

## Discussion

As is clear from the ongoing COVID-19 pandemic, understanding the molecular mechanisms that drive viral sensing and inflammatory pathogenesis during infection remains key to developing rationalized, host-directed treatments to support antiviral defense or quell severe disease. CARD8 is expressed in airway epithelia (*10, 39, 40*) and other cell types, including monocytes and T cells (*26, 28, 41*) that are physiologically relevant for respiratory coronaviruses and picornaviruses (*39, 40, 42*), including SARS-CoV-2 (*29–31*). Our data showing that SARS-CoV-2 infection activates the CARD8 inflammasome in THP-1 cells supports findings that inflammasome activation contributes to severe COVID-19, and suggests that the CARD8 inflammasome in monocytes and macrophages contribute to inflammation in COVID-19 patients.

Based on our results and the finding that FcγR-mediated uptake of SARS-CoV-2 virions into monocytes leads to abortive infection (*31*), we speculate that CARD8-dependent pyroptosis contributes to the poor permissiveness of myeloid cells for SARS-CoV-2 (*43*), wherein the myeloid compartment may not substantially contribute to viral load but likely impacts immunopathology via inflammasome-driven inflammation (*44, 45*). Prior reports have proposed a similar role for the NLRP3 inflammasome in myeloid cells (*29–31*). We favor a unifying model in which CARD8-dependent GSDMD pore formation contributes to NLRP3 inflammasome activation, which also offers an explanation for CARD8-dependent release of IL-1β (*46*). Interestingly, the SARS-CoV-2 3CL^pro^ also cleaves and activates NLRP1 in airway epithelia (*9*). This results in cell death via a non-canonical NLRP1>CASP8>CASP3>GSDME inflammasome pathway, suggesting that the cellular context of inflammasome responses may uniquely shape antiviral defense and/or inflammation. We also note that seasonal coronaviruses are capable of activating CARD8, and we speculate that a multitude of variables such as cell tropism shape the outcome of virus-induced inflammasome activation in antiviral immunity and pathogenesis.

Nevertheless, given the impact of SNPs on human CARD8 sensing of pathogenic viruses, it is tempting to speculate that diminished CARD8 inflammasome activation may be a contributing factor to variation in COVID-19 disease outcomes, and more generally for other human pathogenic coronavirus and picornavirus infections. Further studies are required to establish this connection.

Taken together, our findings establish CARD8 as a rapidly evolving, polymorphic, innate immune sensor of infection by positive-sense RNA viruses. We demonstrate that CARD8 has the capacity to detect viral proteases from at least three viral families that include important human pathogens: *Coronaviridae, Picornaviridae,* and *Retroviridae*. These findings also build on the emerging concept that ETI is an important mechanism of pathogen recognition, including the use of host mimicry of viral polyprotein cleavage motifs as an evolutionary strategy in the ongoing arms race between host and viruses.

## Materials and Methods

### Motif generation and search

To build the betacoronavirus (betaCoV) 3CL^pro^ cleavage motif, 995 nonredundant betaCoV polyprotein sequences were collected from the Viral Pathogen Resource (ViPR) (*47*) and aligned with five well-annotated reference enteroviral polyprotein sequences from RefSeq (**S1 Fig, A and B**). P1 and P1’ of the annotated cleavage sites across the RefSeq sequences served as reference points for putative cleavage sites across the 995 ViPR sequences (**S1 Table**). Four amino acid residues upstream (P4-P1) and downstream (P1’-P4’) of each cleavage site were extracted from every MAFFT-aligned (*48*) polyprotein sequence, resulting in 1000 sets of cleavage sites (RefSeq sites included) (**S1 Table**). Each set of cleavage sites representative of each polyprotein was then concatenated (**S2 Table**). Next, duplicates were removed from the concatenated cleavage sites (**S2 Table**). The remaining 60 nonredundant, concatenated cleavage sites were then split into individual 8-mer cleavage sites and were aligned using MAFFT (*48*) to generate Geneious-defined (*49*) sequence logo information at each aligned position (**S2 Table**). Pseudo-counts to the position-specific scoring matrix were adjusted as described previously (*11*) and a motif p-value cut-off of 0.00231 corresponding to detection of 99% of the initial polyprotein cleavage sites was selected (**S1 FigC**).

### Sequence alignments and phylogenetic trees

Complete polyprotein sequences from 60 betaCoVs with non-redundant cleavage sites (see ‘Motif generation and search’ section above) were downloaded from ViPR. Sequences were aligned using MAFFT (*48*) and a neighbor-joining phylogenetic tree was generated using Geneious software (*49*).

### Evolutionary analyses

For phylogenomic analyses of CARD8 and NLRP1 (**Fig 3A**), human CARD8 (accession NP_001338711) and human NLRP1 (accession NP_127497.1) were used as BLASTP (*50*) search queries against the indicated mammalian proteomes (**Fig 3A**). A species was determined to have an ortholog if it had a protein with >50% sequence identity, >70% sequence coverage, and was the bi-directional best hit to the indicated human protein. The species tree shown in **Fig 3A** is based on NCBI Common Tree (https://www.ncbi.nlm.nih.gov/Taxonomy/CommonTree/wwwcmt.cgi). To identify regions of mammalian CARD8s that are orthologous to the 3CL^pro^ cleavage site (Q349) in *Rousettus aegyptiacus* CARD8, a 50 amino acid region of *R. aegyptiacus* CARD8 that was centered on the Q349 cleavage site was used as a BLASTP query against the entire RefSeq protein database with a 60% sequence identity cut-off. A single CARD8 sequence from each species (**S8 Fig)** was aligned using MAFFT (*48*) and trimmed to only include the eight amino acid spanning the cleavage site.

For positive selection analyses, primate nucleotide sequences that aligned to human full length CARD8 (Protein: NP_001338711, mRNA: NM_001351782.2) were downloaded from NCBI and aligned using MAFFT (*48*). Only eight other primate sequences, only from hominoids and Old World monkeys, were fully alignable to full length human CARD8 (sequence accessions in table S3). Maximum likelihood (ML) tests were performed with codeml in the PAML software suite (*51*) or using BUSTED (*52*) on the DataMonkey (*53*) server. For PAML, aligned sequences were subjected to ML tests using NS sites models disallowing (M7) or allowing (M8) positive selection. The p-value reported is the result of a chi-squared test on twice the difference of the log likelihood (lnL) values between the two models using two degrees of freedom. We confirmed convergence of lnL values by performing each analysis using two starting omega (dN/dS) values (0.4 and 1.5). Results are reported from analyses using the F61 codon frequency model. Analyses with the F3x4 model gave similar results. For evolutionary analyses of regions of CARD8, the full-length alignment was truncated to only include codons 1-161 (‘tripwire’ region) or 162-537 (FIIND-CARD region) and PAML or BUSTED analyses were performed as described above.

We used three independent methods to estimate individual codons within CARD8 that have been subject to positive selection (**S4 Table**). PAML was used to identify positively selected codons with a posterior probability greater than 0.90 using a Bayes Empirical Bayes (BEB) analysis and the F61 or F3x4 codon frequency models. The same CARD8 alignment was also used as input for FEL (*54*) and FUBAR (*55*) using the DataMonkey (*53*) server. In both cases, default parameters were used and codons with a signature of positive selection with a p-value of <0.1 are reported. In all cases, codon numbers correspond to the amino acid position and residue in human CARD8 (NCBI accession NP_001338711).

### Plasmids and constructs

Megabat CARD8, CASP1, IL-1β, and all 3CL^pro^ sequences were ordered as either gBlocks (Integrated DNA Technologies, San Diego, CA) or Twist Gene Fragments (Twist Biosciences, South San Francisco, CA). All sequences are found in **S7 Table**. Vectors containing the coding sequences of human CARD8, ASC, human CASP1, human IL-1β-V5, and TEV^pro^ were previously described (*24*). Vector psPAX2 containing the untagged coding sequence for HIV-1 gag-pol was a gift from Didier Trono (Addgene plasmid # 12260).

For CARD8 cleavage assays, the coding sequences of human CARD8 (NCBI accession NP_001171829.1), human CARD8 mutants (Q37A, Q37A Q61A, S39P, R40W), human CARD8 TEV, *Rousettus aegyptiacus* (megabat) CARD8 (NCBI accession XP_016010896), and megabat CARD8 TEV were cloned into the pcDNA5/FRT/TO backbone (Invitrogen, Carlsbad, CA) with an N-terminal 3xFlag-mCherry tag. For CARD8 activation, the same sequences were cloned into the pQCXIP vector backbone (Takara Bio, Mountain View, CA) with a C-terminal Myc tag. Megabat CASP1 (NCBI accession KAF6464288) and megabat IL-1β (NCBI accession KAF6447073), also from *Rousettus aegyptiacus*, were cloned in the same vector as their respective human orthologues. 3CL^pro^ sequences were cloned with an N-terminal HA tag into the QCXIP vector backbone, flanked by polyprotein cleavage sites fused to N-terminal eGFP and C-terminal mCherry (**S2 Fig)**. 3C^pro^ constructs were described previously (*11*).

Single point mutations were made using overlapping stitch PCR. All plasmid stocks were sequenced across the entire inserted region to verify that no mutations were introduced during the cloning process. The primers used for cloning are described in **S7 Table**.

### Cell culture and transient transfection

All cell lines (HEK293T, HEK-Blue-IL-1β) are routinely tested for mycoplasma by PCR kit (ATCC, Manassas, VA) and kept a low passage number to maintain less than one year since purchase, acquisition or generation. HEK293T cells were obtained from ATCC (catalog # CRL-3216) and HEK-Blue-IL-1β cells were obtained from Invivogen (catalog # hkb-il1b) and all lines were verified by those sources, and were grown in complete media containing DMEM (Gibco, Carlsbad, CA), 10% FBS, and appropriate antibiotics (Gibco, Carlsbad, CA). THP-1 cells were purchased from ATCC, and grown in complete media containing RPMI (Gibco, Carlsbad, CA) 10% FBS, and 1% L-glutamine. For transient transfections, HEK293T cells were seeded the day prior to transfection in a 24-well plate (Genesee, El Cajon, CA) with 500 µl complete media. Cells were transiently transfected with 500 ng of total DNA and 1.5 µl of Transit X2 (Mirus Bio, Madison, WI) following the manufacturer’s protocol. HEK-Blue IL-1β reporter cells (Invivogen, San Diego, CA) were grown and assayed in 96-well plates (Genesee, El Cajon, CA).

### Generation of knockout and transgenic cell lines

*CARD8* knockouts in HEK293T cells were generated similarly to *NLRP1* knockouts described in (*11*). Briefly, lentivirus-like particles were made by transfecting HEK293T cells with the plasmids psPAX2 (gift from Didier Trono, Addgene plasmid # 12260), pMD2.G (gift from Didier Trono, Addgene plasmid # 12259), and either pLB-Cas9 (gift from Feng Zhang, Addgene plasmid # 52962) (*56*) or plentiGuide-Puro, which was adapted for ligation-independent cloning (gift from Moritz Gaidt) (*57*). Conditioned supernatant was harvested 48 and 72 hours post-transfection and used for spinfection of HEK293T cells at 1200 x *g* for 90 minutes at 32°C. Forty-eight hours post-spinfection, cells with stable expression of Cas9 were selected in media containing 100 µg/ml blasticidin. Blasticidin-resistant cells were then transduced with sgRNA-encoding lentivirus-like particles, and selected in media containing 0.5 µg/ml puromycin. Cells resistant to blasticidin and puromycin were single cell cloned by limiting dilution in 96-well plates, and confirmed as knockouts by Sanger sequencing. *CARD8* knockout THP-1 cells were generated as described previously (*58*). Briefly, a CARD8 specific sgRNA was designed using CHOPCHOP (*59*), and cloned into a plasmid containing U6-sgRNA-CMV-mCherry-T2A-Cas9 using ligation-independent cloning. THP-1 cells were electroporated using the BioRad GenePulser Xcell. After 24 h, mCherry-positive cells were sorted and plated for cloning by limiting dilution. Monoclonal lines were validated as knockouts by deep sequencing and OutKnocker analysis, as described previously (*60, 61*). Knockout lines were further validated by immunoblot and functional assays. sgRNA used to generate knockouts are described in **S7 Table**. To make THP-1 cells susceptible to SARS-CoV-2 infection(*62*), ACE2 and TMPRSS2 expressing THP-1 cells were made using the same lentiviral transduction protocol as described above, but using the transfer plasmid PpWIP-IRES-Bla-AK-ACE2-IRES-TMPRSS2 (gift from Sonja Best). THP-1 cells were selected with 10 µg/ml blasticidin.

### THP-1 treatments and viral infections

50,000-100,000 THP-1 cells per well in 96-well round bottom plates in 50 µl OptiMEM containing 500 ng/mL Pam3CSK4 for 6h, followed by treatment with Val-boroPro (10µM) or infection with the coronaviruses hCoV-229E (BEI NR-52726) or SARS-CoV-2 (USA/WA-1/2020, a gift from Dr. Ralph Baric) at indicated multiplicities of infection (MOIs). 48h post-treatment or infection, supernatants were harvested for the detection of IL-1β (see below). Cells were transferred to a white-walled 96-well assay plate and mixed with an equal volume of Cell Titer Glo reagent (Promega). Measurements for fluorescence at 544-15 nm (excitation), 620-20 nm (emission) or luminescence at 555-70 (emission) were taken following incubation at room temperature, rocking for 10min.

### CARD8 cleavage assays

100 ng of epitope-tagged human CARD8 (WT, Q37A, Q37A Q61A, S39P, R40W), human CARD8 TEV, megabat CARD8 WT or megabat CARD8 TEV was co-transfected with either HA-tagged QCXIP empty vector (‘-’), 250 ng of TEV^pro^, 250 ng of untagged HIV-1^pro^ (HIV-1 gag-pol carrying HIV-1 protease activity), 5 ng of HA-tagged 3CL^pro^, or 250 ng of HA-tagged 3C^pro^-encoding constructs. Twenty-four hours post-transfection, the cells were harvested, lysed in 1x NuPAGE LDS sample buffer (Invitrogen, Carlsbad, CA) containing 5% β-mercaptoethanol (Fisher Scientific, Pittsburg, PA) and immunoblotted with antibodies described in **S8 Table**.

### CARD8 activity assays

To reconstitute the human CARD8 inflammasome, 100 ng of human CASP1 and 50 ng of human IL-1β-V5 were co-transfected with 50 ng of either HA-tagged QCXIP empty vector, wild-type or mutant pQCXIP-CARD8-Myc constructs in *CARD8* KO HEK293T cells. To reconstitute the megabat CARD8 inflammasome, 10 ng of megabat CASP1 and 50 ng of megabat IL-1β-V5 were co-transfected with 2 ng megabat CARD8 constructs in *CARD8* KO HEK293T cells. These cells were further co-transfected with either empty vector (‘-’), 250 ng of TEV^pro^, 100 ng of untagged HIV-1 gag-pol (with HIV-1 protease activity), 5 ng of HA-tagged 3CL^pro^, 100 ng of HA-tagged enteroviral 3C^pro^, or 20 ng of HA-tagged non-enteroviral 3C^pro^-encoding constructs. Twenty-four hours post-transfection, cells were harvested and lysed in 1x NuPAGE LDS sample buffer containing 5% β-mercaptoethanol and immunoblotted with antibodies described in **S8 Table** or culture media was harvested for quantification of IL-1β levels by HEK-Blue assays (see below). Appearance of the mature p17 band of IL-1β indicates successful assembly and activation of the inflammasome.

### HEK-Blue IL-1β assay

To quantify the levels of bioactive IL-1β released from cells, we employed HEK-Blue IL-1β reporter cells (Invivogen, San Diego, CA). In these cells, binding to IL-1β to the surface receptor IL-1R1 results in the downstream activation of NF-kB and subsequent production of secreted embryonic alkaline phosphatase (SEAP) in a dose-dependent manner (*11*). SEAP levels are detected using a colorimetric substrate assay, QUANTI-Blue (Invivogen, San Diego, CA) by measuring an increase in absorbance at OD655.

Culture supernatant from inflammasome-reconstituted HEK293T cells or HEK293T *CARD8* KO cells that had been transfected with 3CL pro was added to HEK-Blue IL-1β reporter cells plated in 96-well format in a total volume of 200 µl per well. On the same plate, serial dilutions of recombinant human IL-1β (Invivogen, San Diego, CA) were added to generate a standard curve for each assay. Twenty-four hours later, SEAP levels were assayed by taking 20 µl of the supernatant from HEK-Blue IL-1β reporter cells and adding to 180 µl of QUANTI-Blue colorimetric substrate following the manufacturer’s protocol. After incubation at 37°C for 30–60 min, absorbance at OD655 was measured on a BioTek Cytation five plate reader (BioTek Instruments, Winooski, VT) and absolute levels of IL-1β were calculated relative to the standard curve. All assays, beginning with independent transfections or infections, were performed in triplicate.

### Immunoblotting and antibodies

Harvested cell pellets were washed with 1X PBS, and lysed with 1x NuPAGE LDS sample buffer containing 5% β-mercaptoethanol at 98C for 10 min. The lysed samples were spun down at 15000 RPM for two minutes, followed by loading into a 4–12% Bis-Tris SDS-PAGE gel (Life Technologies, San Diego, CA) with 1X MOPS buffer (Life Technologies, San Diego, CA) and wet transfer onto a nitrocellulose membrane (Life Technologies, San Diego, CA). Membranes were blocked with PBS-T containing 5% bovine serum albumin (BSA) (Spectrum, New Brunswick, NJ), followed by incubation with primary antibodies for V5 (IL-1β), FLAG (mCherry-fused CARD8 for protease assays), Myc (CARD8-Myc for activation assays), HA (viral protease), or GAPDH. Membranes were rinsed three times in PBS-T then incubated with the appropriate HRP-conjugated secondary antibodies. Membranes were rinsed again three times in PBS-T and developed with SuperSignal West Pico PLUS Chemiluminescent Substrate (Thermo Fisher Scientific, Carlsbad, CA). The specifications, source, and clone info for antibodies are described in **S8 Table**.

## Acknowledgments

We thank all members of the Daugherty and Mitchell laboratories and Michael Emerman for helpful discussions. We thank Joshua Marceau, Nell Baumgarten, and Julie Overbaugh for providing HCoV-229E resources and expertise.

## Funding

National Institutes of Health grant R35 GM133633 (MDD)

Pew Biomedical Scholars Program (MDD)

Hellman Fellows Program (MDD)

Burroughs Wellcome Investigators in the Pathogenesis of Infectious Disease Program (MDD)

National Institutes of Health T32 grant GM007240 (BVT, APR, CB, LKC)

National Science Foundation graduate research fellowship 2019284620 (CB)

National Institutes of Health grant DP2 AI 154432-01 (PSM)

Mallinckrodt Foundation grant (PSM)

National Institutes of Health T32 grant GM007270 (JK)

UC Berkeley CEND Catalyst award (REV)

National Institutes of Health grant R37AI075039 (REV)

Helen Hay Whitney Postdoctoral Fellowship (NSG)

Investigator of the Howard Hughes Medical Institute (REV)

## Author contributions

Conceptualization: BVT, PSM, MDD

Methodology: BVT, NSG, JK, APR, EAT, PSM, MDD

Investigation: BVT, RA, NSG, JK, APR, LKC, CMB, EAT, EJF, PSM, MDD

Visualization: BVT, NSG, PSM, MDD

Funding acquisition: REV, PSM, MDD

Project administration: REV, JLH, RS, PSM, MDD

Supervision: REV, JLH, RS, PSM, MDD

Writing – original draft: BVT, PSM, MDD

Writing – review & editing: All authors

## Supporting Information

**S1 Fig.**
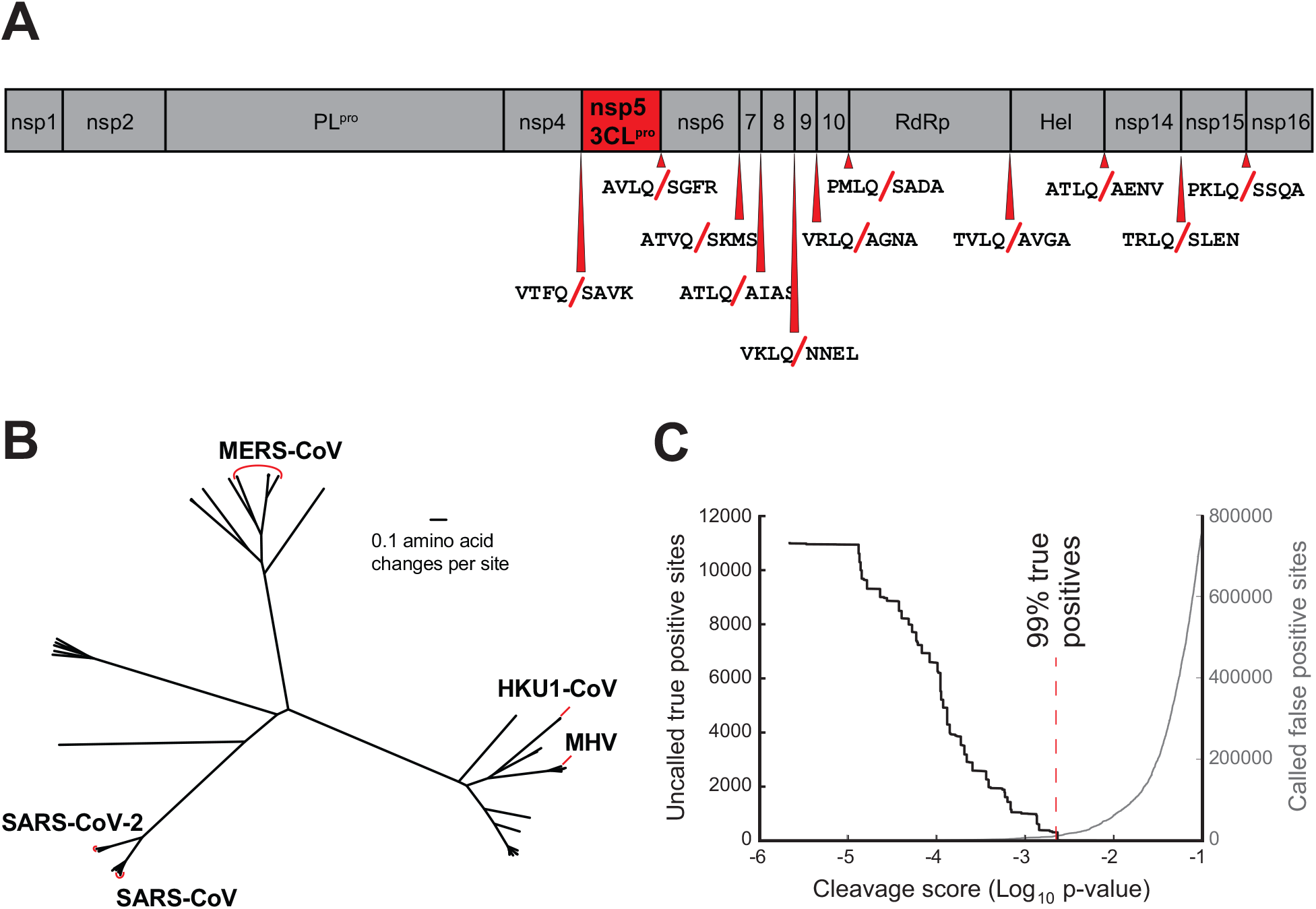
Motif generation of coronaviral 3CL^pro^ polyprotein cleavage. (A) Schematic of 3CL^pro^ cleavage sites (11 red triangles) within the polyprotein of SARS-CoV-2 (SARS-2), the causative agent of COVID-19. Shown are four amino acids flanking each side of the cleavage site within the polyprotein. (B) Phylogenetic tree of 60 coronavirus polyprotein coding sequences depicting the betacoronaviruses sampled in this study with human relevant coronaviruses labeled (table S2). (C) Training set data used to determine the motif search threshold for FIMO (table S1). The X-axis represents a log10 of the p-value reported by FIMO as an indicator for the strength of the cleavage motif hit (cleavage score). (Left) The Y-axis depicts the number of uncalled true positives, or motif hits that overlap with the initial set of 8mer polyprotein cleavage sites used to generate the motif, in the training set of coronavirus polyprotein sequences (black line). (Right) The Y-axis depicts the number of called false positive sites, or any motif hits found in the polyprotein that are not known to be cleaved by 3CL^pro^, in the training set of coronaviral polyprotein sequences (gray). A red vertical dashed line indicates the threshold that captures 99% of true positive polyprotein hits (p-value = 0.00231).

**S2 Fig.**
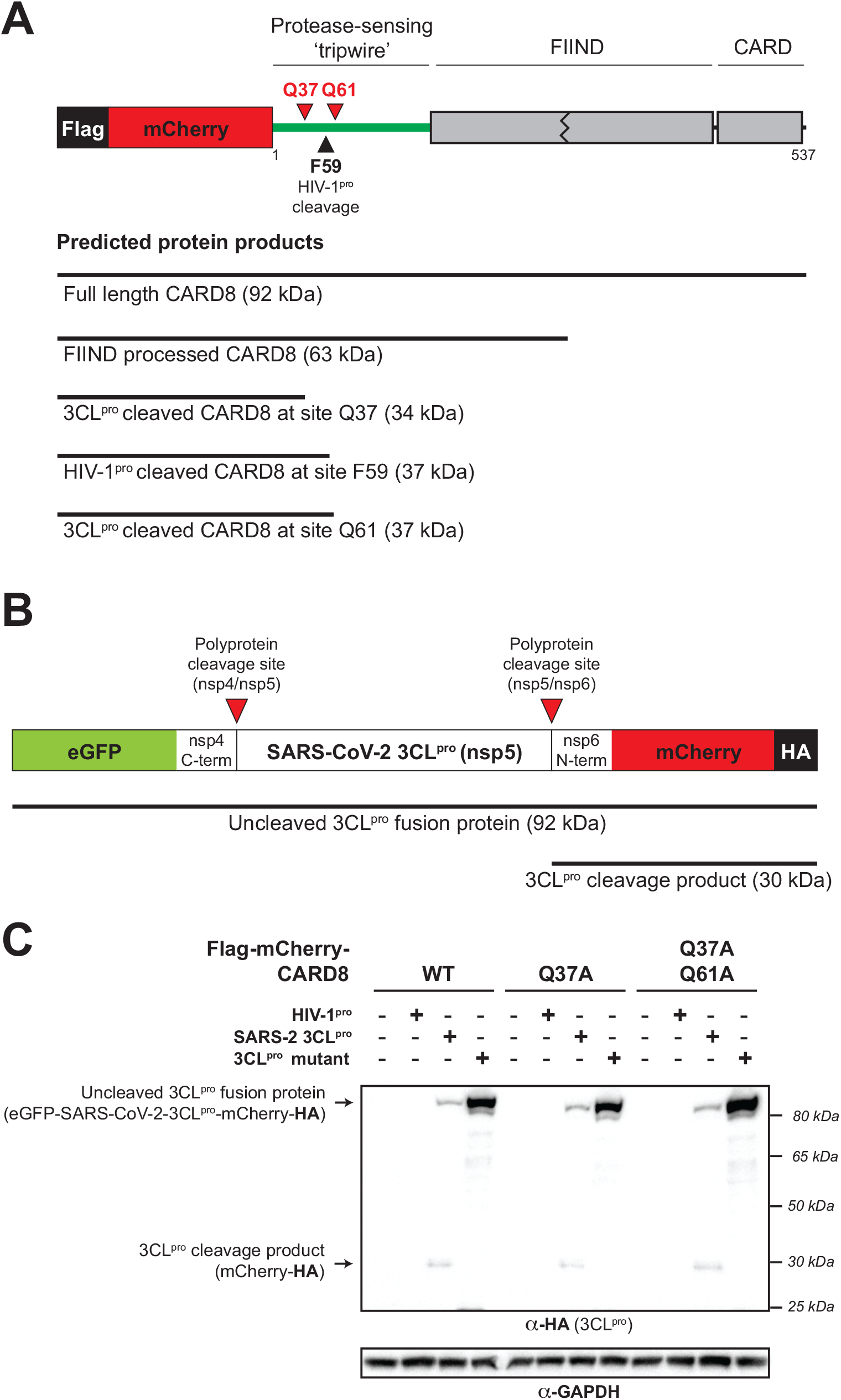
Schematic of CARD8 and 3CL^pro^ expression constructs for cleavage assays and immunoblot depicting HA-tagged SARS-2 3CL^pro^ and SARS-2 3CL^pro^ mutant. (A) Schematic of the expression constructs used for CARD8 cleavage assays. Full length CARD8 was fused to a 3xFlag-mCherry domain to increase the stability and our ability to detect viral protease cleavage products in the N-terminus of CARD8. Below are shown the expected sizes of full length and FIIND processed CARD8, as well as the expected sizes that would result from viral protease cleavage at the indicated sites. (B) Schematic of the expression constructs used for 3CL^pros^ (nsp5s). A region of the viral polyprotein spanning the C-terminal nine residues of nsp4 through the N-terminal nine residues of nsp6 was cloned between eGFP and mCherry-HA. Inactive protease is expressed as a full-length fusion protein (92 kDa predicted molecular weight). Active 3CL^pro^ cleaves at the indicated polyprotein cleavage sites (red triangles), liberating the active protease from the construct and resulting in an HA-tagged mCherry product (30 kDa predicted molecular weight). (C) An expanded view of the anti-HA-stained immunoblot shown in Figure 1B highlighting uncleaved (Uncleaved 3CL^pro^ fusion protein) and cleaved (3CL^pro^ cleavage product) HA-tagged protein products.

**S3 Fig.**
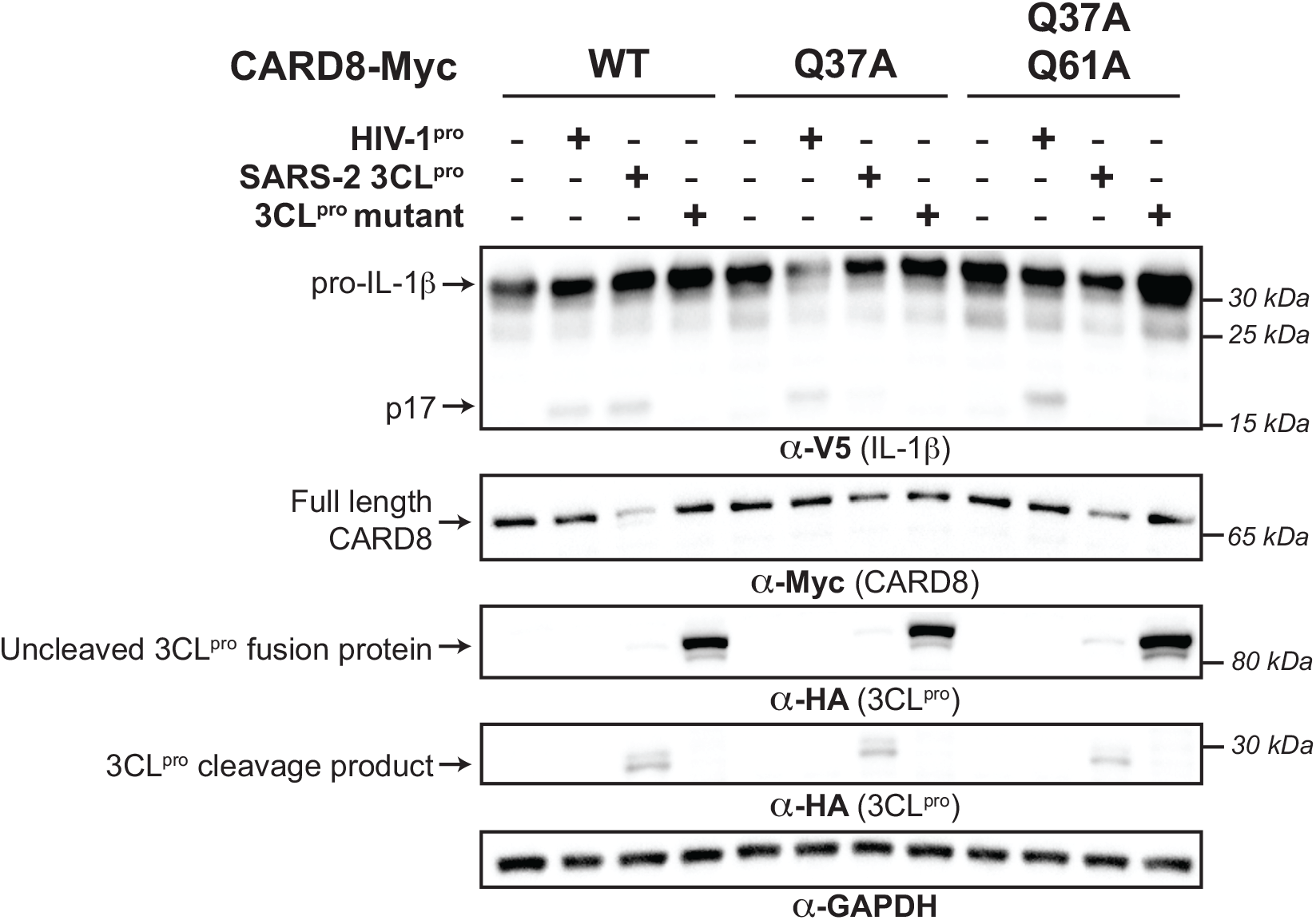
CARD8 cleavage by viral proteases results in inflammasome activation and IL-1β maturation. CARD8 inflammasome assay. CARD8 KO HEK293T cells were co-transfected using the indicated Myc-tagged CARD8 plasmid constructs, V5-IL-1β, CASP1, and HA-tagged protease constructs (SARS-CoV-2 3CL^pro^ (SARS-2 3CL^pro^), SARS-CoV-2 3CL^pro^ catalytic mutant C145A (3CL^pro^ mutant), HIV-1 gag-pol (HIV-1^pro^), or empty vector (-)). Appearance of a mature bioactive IL-1β (p17) indicates inflammasome activation.

**S4 Fig.**
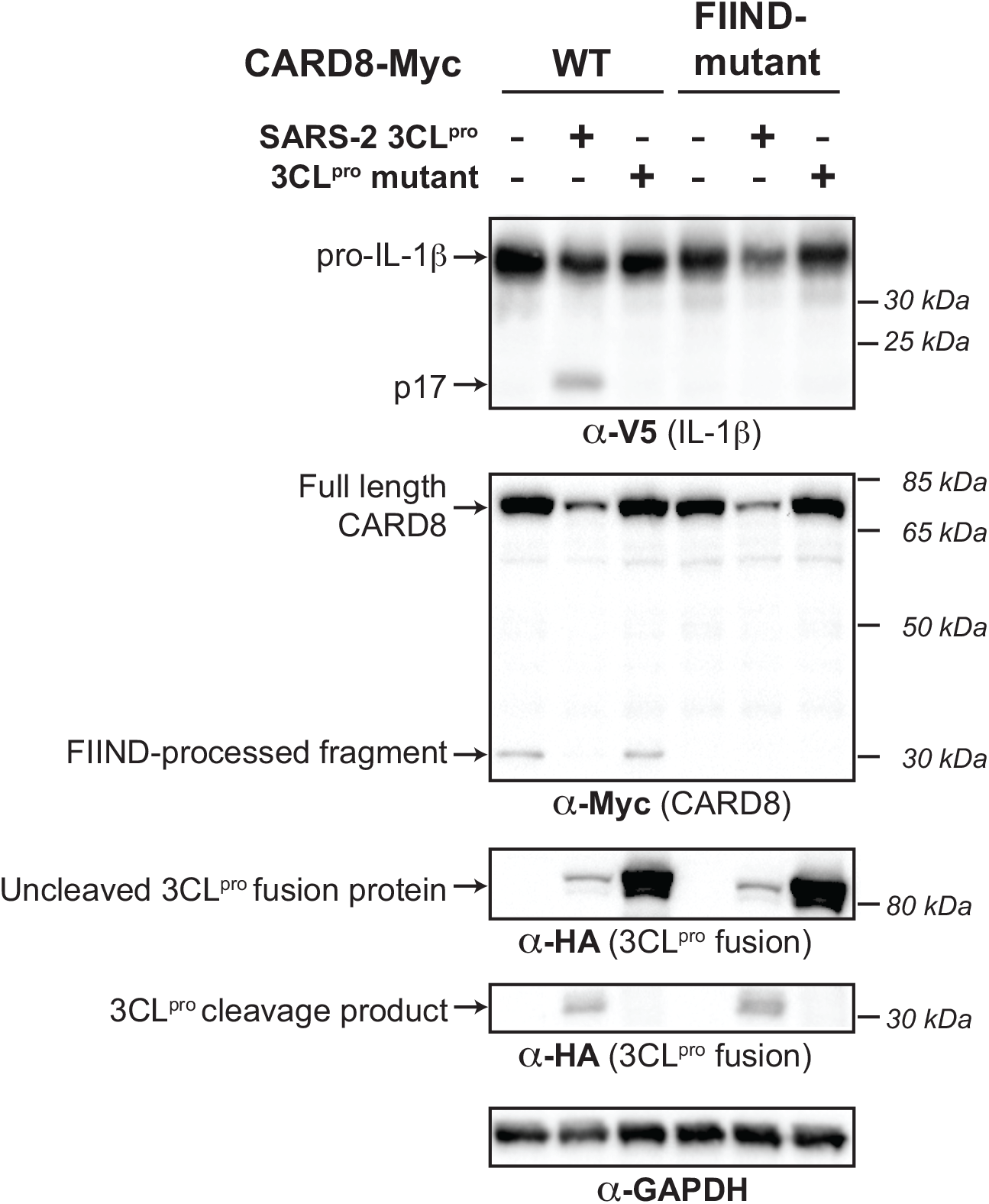
3CLpro-mediated activation of the human CARD8 inflammasome depends on FIIND autoprocessing. CARD8 inflammasome activation assay depicting loss of CARD8 activation with a FIIND autoprocessing mutant (S297A).

**S5 Fig.**
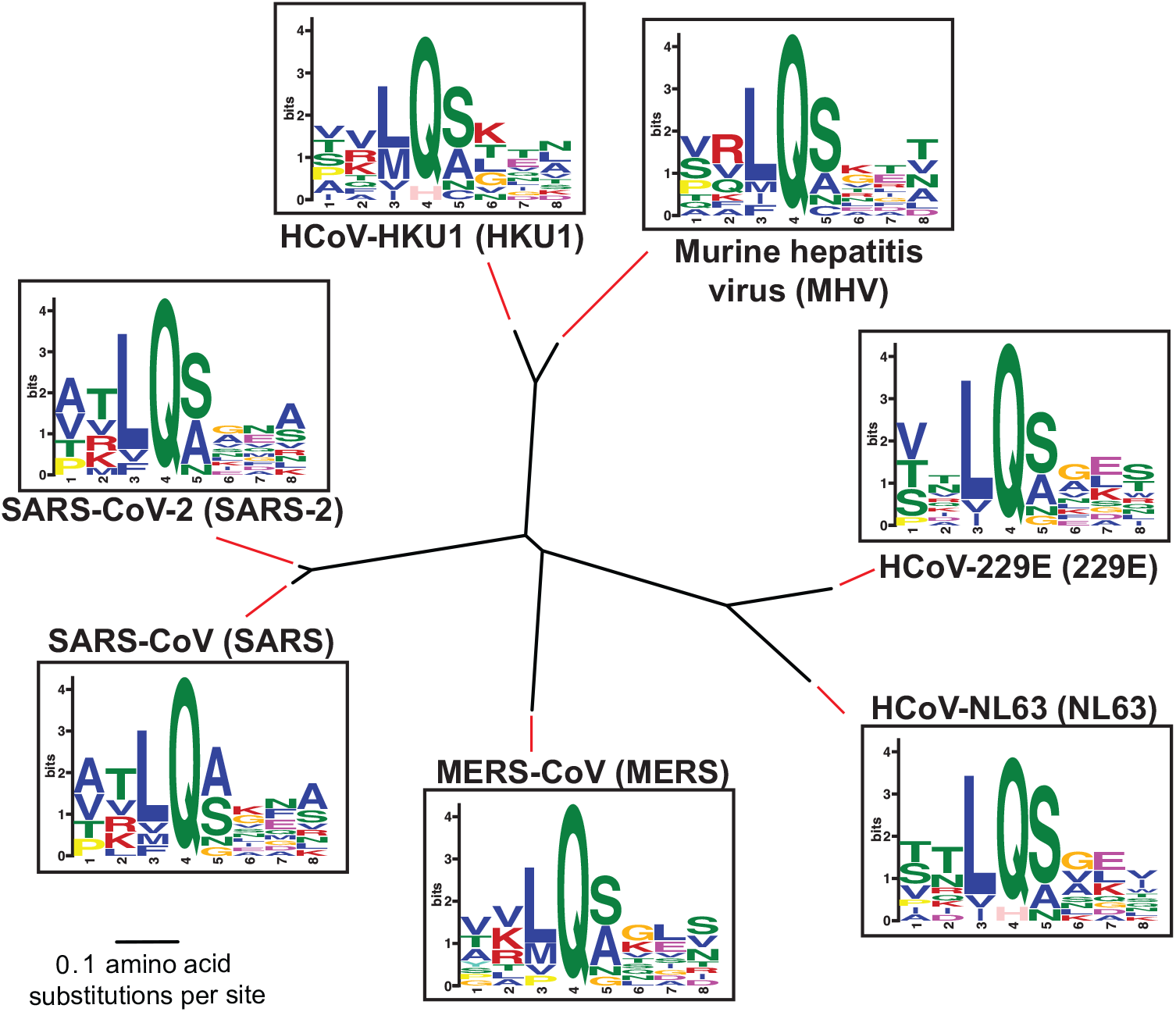
3CLpros used in this study demonstrate similar polyprotein cleavage. Phylogenetic tree of 3CL^pro^ protein sequences used in this study from the indicated coronaviruses. Shown next to the virus name is the sequence motif generated from the 3CL^pro^ polyprotein cleavage sites from that specific virus.

**S6 Fig.**
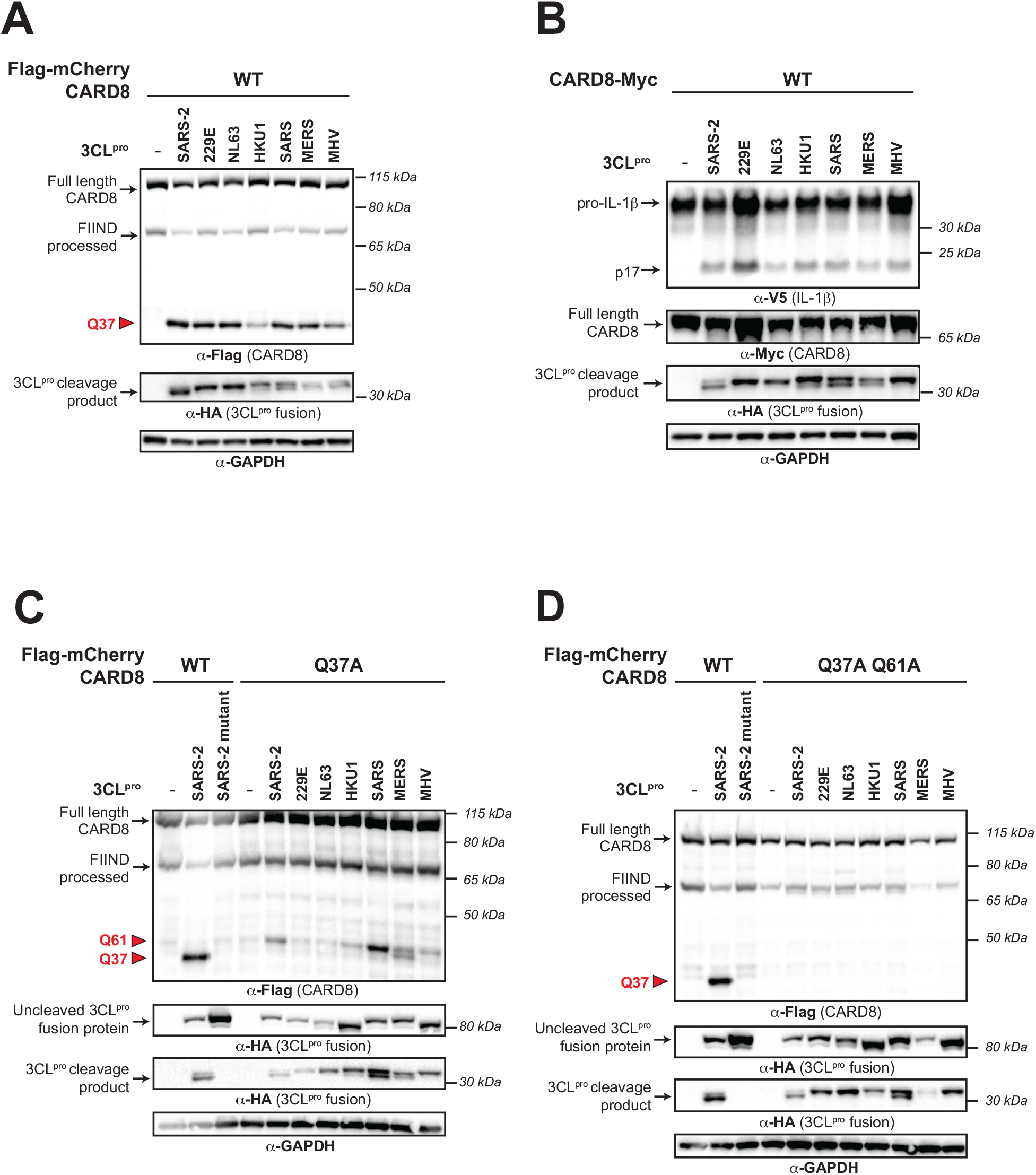
Site-specific CARD8 cleavage and inflammasome activation by diverse coronavirus 3CL^pros^. (A) Cleavage assay depicting cleavage of human CARD8 by the indicated 3CL^pros^ from diverse coronaviruses (SARS-CoV-2 (SARS-2), HCoV-229E (229E), HCoV-NL63 (NL63), HCoV-HKU1 (HKU1), SARS-CoV (SARS), MERS-CoV (MERS), and murine hepatitis virus (MHV)). (B) CARD8 inflammasome activation assay with human CARD8 WT and the indicated 3CL^pro^. (C-D) Cleavage assays mapping the cleavage specificity of diverse 3CL^pros^. Indicated proteases were co-transfected with WT and Q37A (C) or Q37A Q61A (D).

**S7 Fig.**
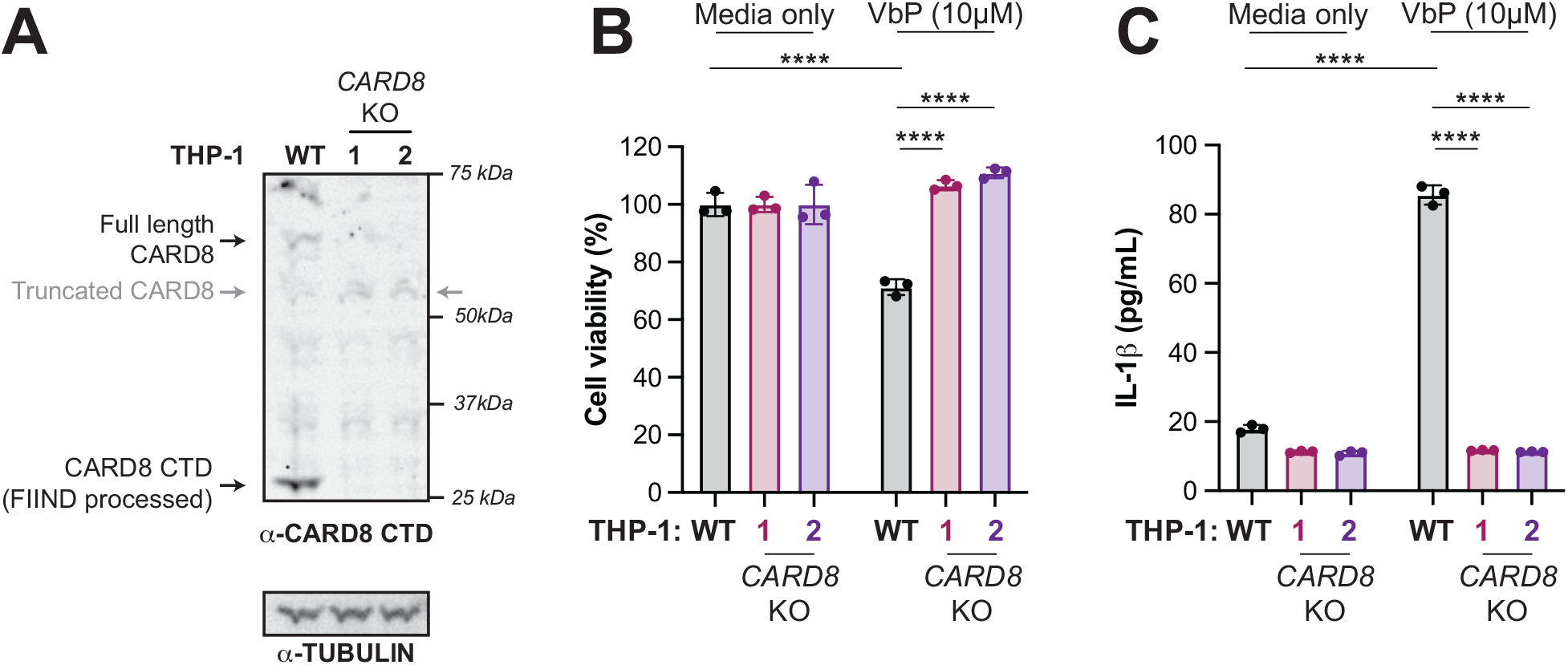
Validation of *CARD8* KO THP-1 cells. (A) Immunoblot of wildtype (WT) or *CARD8* knockout (KO) 1 and KO2 THP-1 cells. The sgRNA used to edit *CARD8* results in a truncated CARD8 (grey arrow) that removes the C-terminal domain (CTD), including the CARD. (B-C) Indicated THP-1 cells were primed with 0.5 µg/mL Pam3CSK4 for 6h, followed by treatment with 10 µM Val-boroPro (VbP) or media only. Cell viability (B) was measured 48h post-treatment via the Cell Titer Glo assay and IL-1β levels (C) were measured from cell supernatants using the IL1R reporter assay as in Fig 1D (see **Methods**). Data presented are representative of experiments performed in triplicate. Data were analyzed using two-way ANOVA with Šidák’s post-test: p<0.0001.

**S8 Fig.**
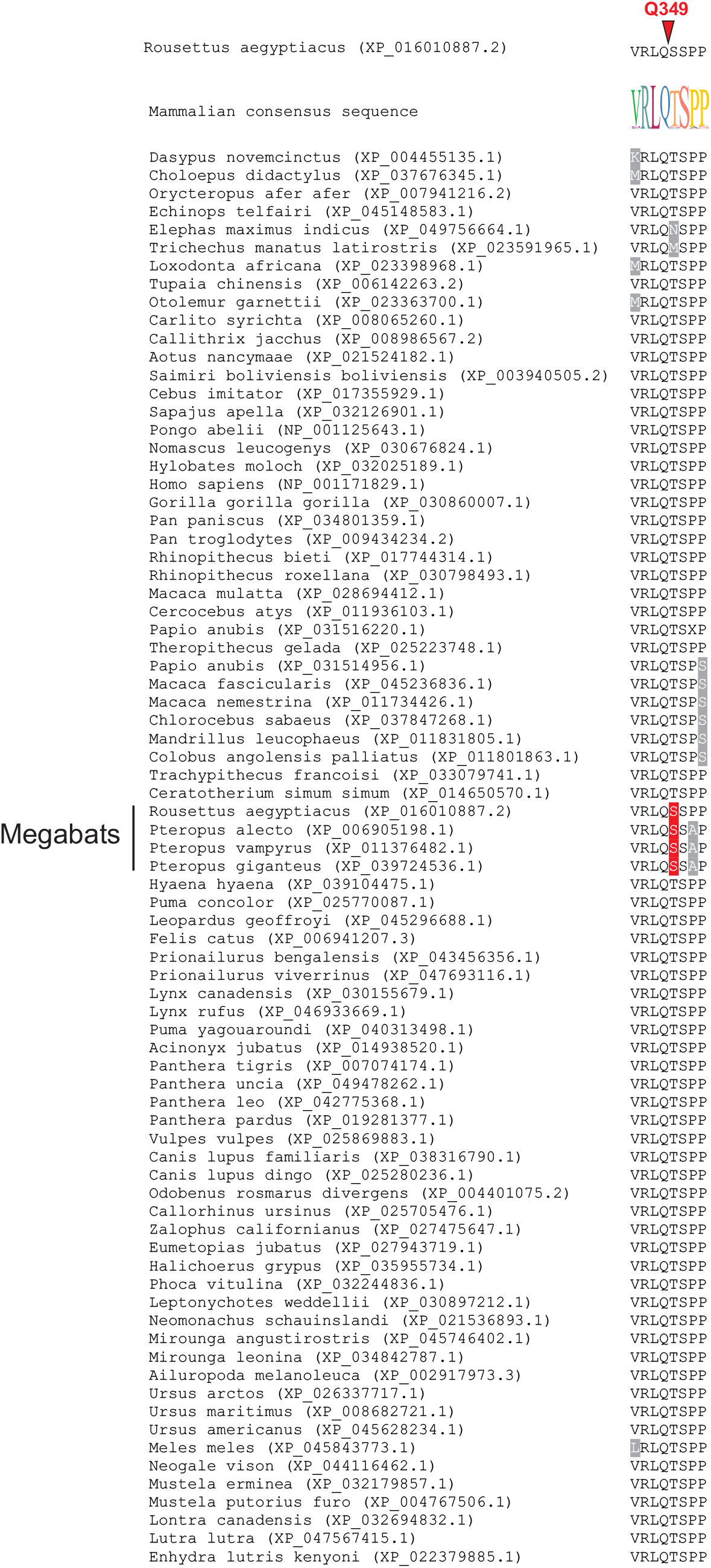
A coronavirus 3CL^pro^ cleavage site in the CARD8 C-terminus is unique to megabats. An alignment of protein sequences homologous to the coronavirus 3CL^pro^ cleavage site in megabat (*Rousettus aegyptiacus*) CARD8 is shown for indicated species (scientific name, accession number). Amino acid numbering is based on *R. aegyptiacus* CARD8. Changes relative to the consensus, depicted as a sequence logo at the top of the alignment) are highlighted. Four species of megabats are indicated, and are unique in having a serine in the P1’ position of the cleavage site (highlighted in red). Most other species have a threonine in this position, which makes the protein uncleavable at this site (Fig 3E).

**S9 Fig.**
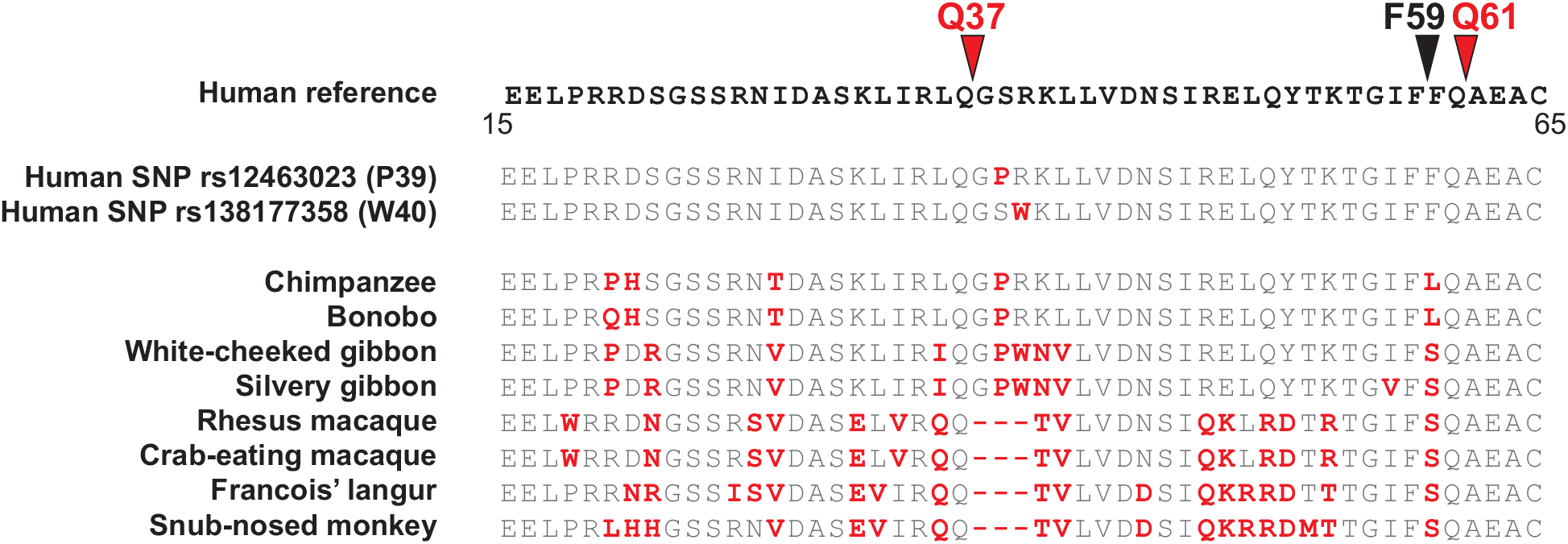
Rapid evolution and human polymorphism in the ‘tripwire’ region of CARD8 that is targeted by viral proteases. An alignment of the CARD8 N-terminus (amino acids 15-65, where amino acid numbering is based on human CARD8) or ‘tripwire’ region from humans and selected non-human primates. Human non-synonymous single nucleotide polymorphism (SNP) that encode CARD8 P39 and W40 are shown. Differences relative to the human reference protein sequence (accession NP_001338711) are indicated in red font. Coronavirus 3CL^pro^ (Q37 and Q61; red arrows) and HIV-1^pro^ cleavages sites in CARD8 are shown. ‘-‘ = indel.

**S10 Fig.**
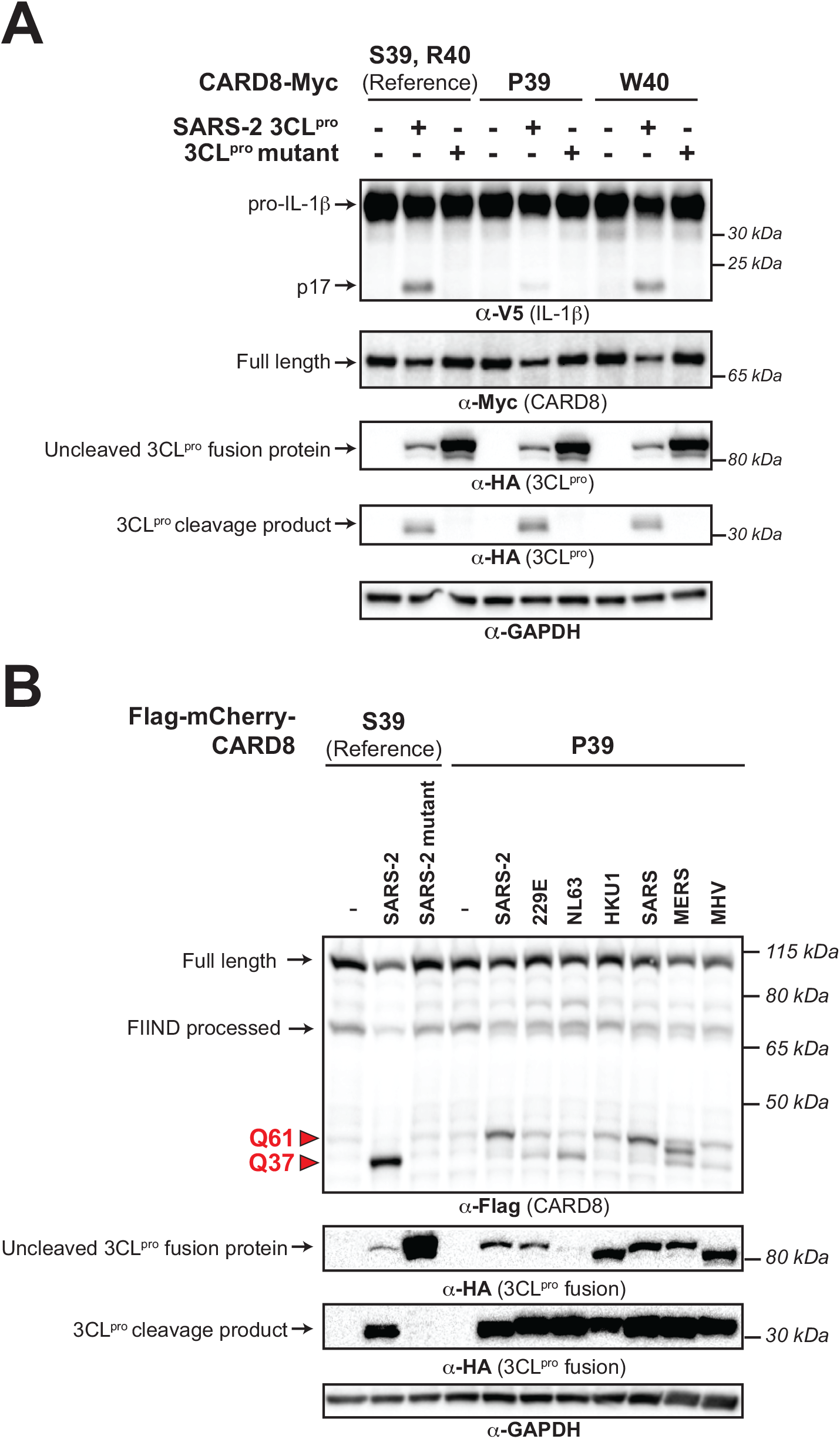
The human CARD8 S39P variant has reduced sensitivity to coronavirus 3CL^pro^ cleavage and inflammasome activation. (A) CARD8 inflammasome activation assay, where inflammasome activation is measured by CASP1-dependent processing of pro-IL-1β to p17. *CARD8* knockout HEK293T cells were co-transfected with Myc-tagged constructs encoding the reference allele of human CARD8 (protein accession NP_001338711, mRNA accession NM_001351782.2) or the human CARD8 non-synonymous single nucleotide polymorphism (SNP) CARD8-P39 (rs12463023) or CARD8-W40 (rs138177358) variants and the HA-tagged SARS-CoV-2 (SARS-2) 3CL^pro^. (B) Comparison of CARD8 S39 and CARD8 P39 cleavage by diverse coronavirus 3CL^pros^. *CARD8* knockout HEK293T cells were co-transfected using the indicated Flag-tagged mCherry-CARD8 fusion constructs with HA-tagged protease constructs (empty vector (‘-’), SARS-2 catalytically inactive mutant (3CL^pro^ mutant) or active 3CL^pro^ from SARS-2, 229E, NL63, HKU1, SARS, MERS, or MHV. Red triangles indicate cleavage sites 3CL^pro^.

**S11 Fig.**
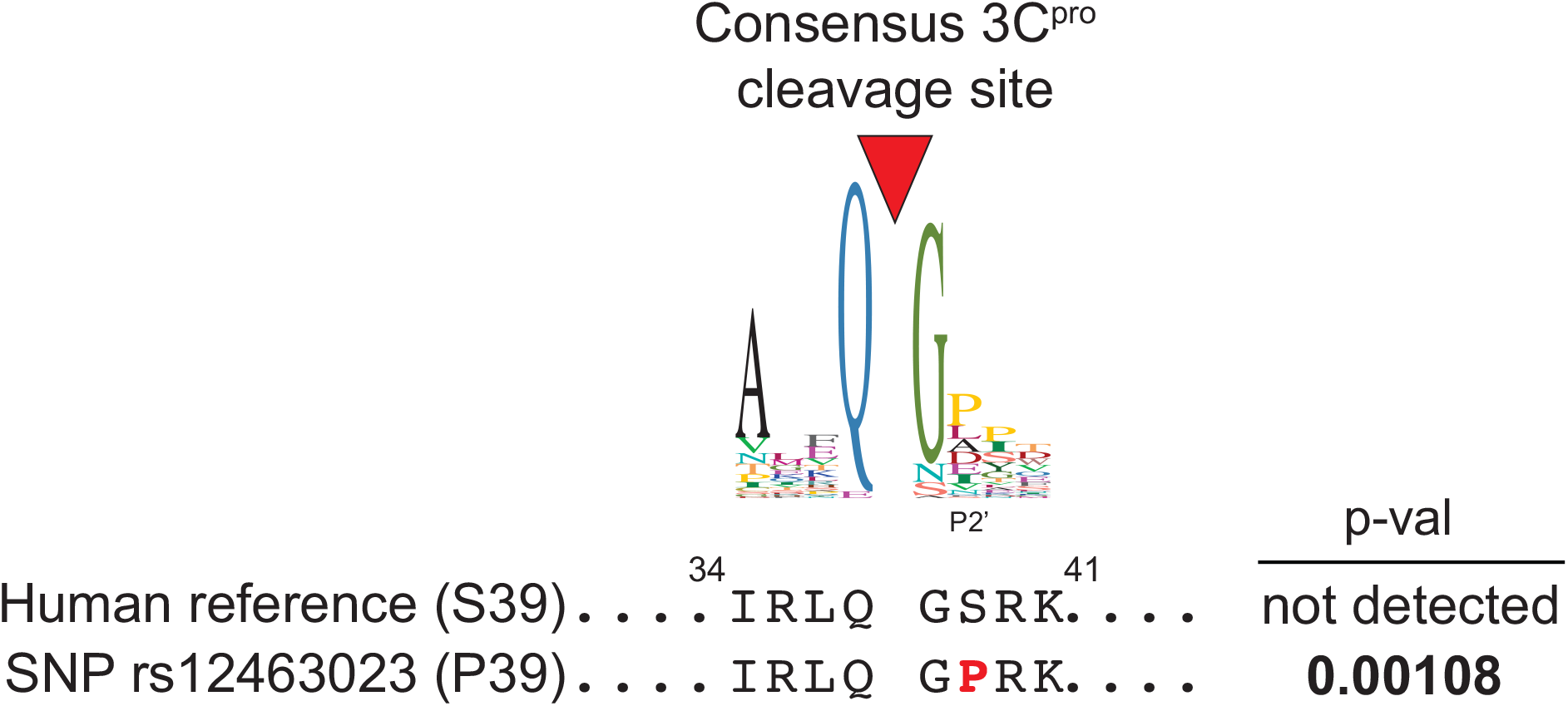
A human SNP is predicted to result in a CARD8 protein that is cleavable by picornavirus 3C^pros^. A previously generated 3C^pro^ consensus cleavage motif for enteroviruses (a genus within picornaviruses), in which a proline is most common in the P2’ position, is shown (*11*). Using this motif to search the human reference CARD8 protein sequence, which contains a serine at residue 39 (S39), does not generate a predicted cleavage site at Q37. However, a human SNP (rs12463023) results in a proline at residue 39 (P39), which generates a predicted 3C^pro^ cleavage site.

**S12 Fig.**
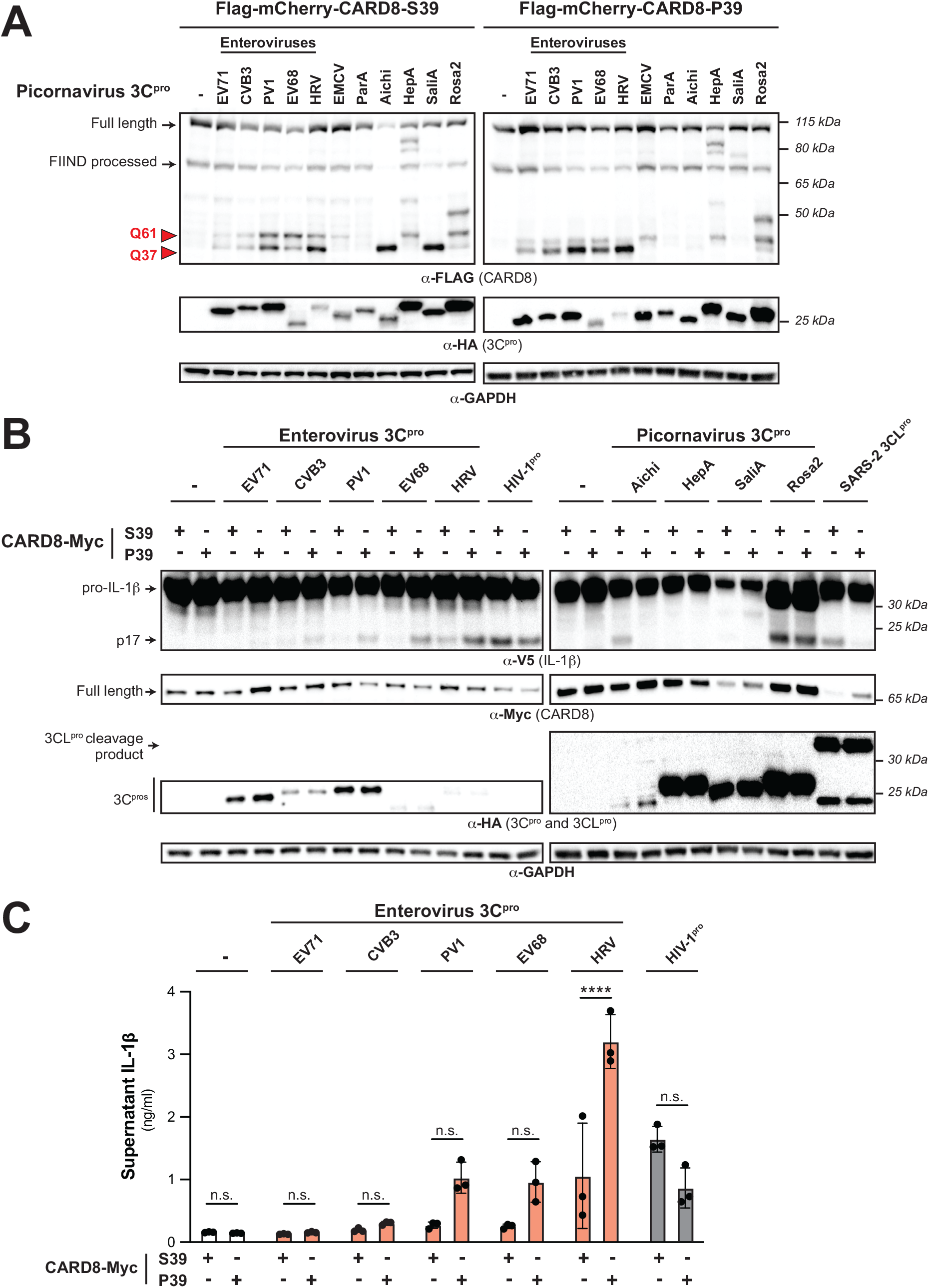
The CARD8 P39 variant is variably susceptible to picornavirus 3C^pros^. (A) WT HEK293T cells were co-transfected using the indicated Flag-tagged mCherry-CARD8 fusion plasmid constructs with V5-IL-1β, CASP1, and HA-tagged 3C^pro^ constructs or empty vector (‘-’). 3C^pros^ from the following viruses were used: enterovirus A71 (EV71), coxsackievirus B3 (CVB3), poliovirus 1 (PV1), enterovirus D68 (EV68), human rhinovirus A (HRV), encephalomyocarditis virus (EMCV), human parechovirus A (ParA), Aichi virus (Aichi), hepatitis A virus (HepA), human salivirus A (SaliA), and human rosavirus 2 (Rosa2). Red arrows denote CARD8 fragments resulting from 3C^pro^ cleavage at indicated sites. (B) CARD8 inflammasome activation assay, where inflammasome activation is measured by CASP1-dependent processing of pro-IL-1β to p17. CARD8 knockout HEK293T cells were co-transfected with Myc-tagged constructs encoding the reference allele of human CARD8 (protein accession NP_001338711, mRNA accession NM_001351782.2) or the human CARD8 non-synonymous single nucleotide polymorphism (SNP) CARD8-P39 (rs12463023) and the indicated 3C^pro^, 3CL^pro^, or HIV-1^pro^ constructs, or empty vector (‘-’). (C) IL-1β assay, which measures the release of bioactive IL-1β in the culture supernatant of cells transfected as described in (B). Individual values (n=3), averages, and standard deviations shown are representative of experiments performed in triplicate. Data were analyzed using one-way ANOVA with Tukey’s post-test. n.s. = not significant. **** = p<0.0001.

**S1 Table. VIPR and RefSeq betacoronaviral polyproteins with 3CL cleavage site concatenations.**

**S2 Table. Betacoronaviral polyproteins with unique 8mer 3CL cleavage site concatenations.**

**S3 Table. Accession numbers of CARD8 sequences used for evolutionarily analyses.**

**S4 Table. Codon positions in full length CARD8 from hominoids and Old World monkeys predicted to be evolving under recurrent positive selection by PAML, FUBAR, and FEL analyses.**

**S5 Table. CARD8 missense variants (>100 allele counts) mapped to GRCh38 reported in gnomAD v3.1.2.**

**S6 Table. CARD8 missense variants (>100 allele counts) mapped to GRCh38 reported in gnomAD v3.1.2 represented by population with the highest allele frequency.**

**S7 Table. Primers, gBlocks, Twist fragments, and sgRNA. S8 Table. List of antibody specifications.**

